# Cloning, heterologous expression and characterization of o-phthalyl-CoA decarboxylase from phthalate degrading denitrifying bacterium

**DOI:** 10.1101/2022.07.29.502009

**Authors:** Madan Junghare

**Affiliations:** Department of Biology, Microbial Ecology, University of Konstanz, Konstanz D-78457, Germany; Faculty of Chemistry, Biotechnology and Food Science, Norwegian University of Life Sciences (NMBU), Ås, Norway

**Keywords:** benzoyl-CoA, decarboxylase, *ortho*-phthalyl-CoA, succinyl-CoA transferase, proteomics.

## Abstract

Phthalic acid esters (phthalates) are used as additives in various plastics and industrial applications. They are produced worldwide in huge amounts causing major pollution in the environment. Biodegradation of phthalates from the environment is an important route for their removal. In our previous work, we showed that *Azoarcus* sp. strain PA01 catabolizes *o*-phthalate via the anaerobic benzoyl-CoA pathway that involved two putative enzymes: the succinyl-CoA:*o*-phthalate CoA-transferase activates *o*-phthalate to *o*-phthalyl-CoA which is subsequently decarboxylated to benzoyl-CoA by *o*-phthalyl-CoA decarboxylase. In this work, we provide the information on the enzymes involved in the promising step of anoxic decarboxylation of *o*-phthalate to benzoyl-CoA. We have identified that there are two proteins are involved in decarboxylation step, of which only one does the actual decarboxylation but other one is essential. *o*-Phthalyl-CoA decarboxylase (PhtDa and PhtDb) encoded by the two genes PA01_00217 and PA01_00218 which catalyses the decarboxylation of activated *o*-phthalate to benzoyl-CoA. Both genes are originally annotated as an UbiD-like/UbiX-like protein. The gene with locus tag PA01_00217 is 1584 bp long coding for protein PhtDa (60 kDa), whereas PA01_00218 is 600 bp long codes for protein PhtDb (22 kDa). Here, we demonstrate that PhtDb is a flavin mononucleotide (FMN)-binding protein which does not function as a decarboxylase alone. Rather, PhtDb is assumed to generate a modified FMN-containing cofactor that is required by the PhtDa for decarboxylase activity. Alone, PhtDa does not function as a decarboxylase either. Recombinantly expressed PhtDa and PhtDb together showed activity for decarboxylation of *o*-phthalyl-CoA to benzoyl-CoA, only if PhtDb was previously incubated with FMN and dimethylallyl monophosphate. Phylogenetically, the proteins PhtDa and PhtDb are closely related to UbiD-like/UbiX-like enzymes that catalyses the decarboxylation of 4-hydroxy-3-octaprenylbenzoic acid to 2-octaprenylphenol, an intermediate step in ubiquinone biosynthesis. Furthermore, multiple sequence alignment and structural modelling of both proteins suggested that only PthDb possesses the binding site for FMN. These results strongly indicate that the flavin-containing cofactor is essential for decarboxylation of *o*-phthalyl-CoA to benzoyl-CoA during anaerobic *o*-phthalate degradation by *Azoarcus* sp. strain PA01.

## Introduction

*o*-Phthalic acid (1,2-dicarboxybenzene) is a synthetic organic compound most used in the manufacturing of phthalic acid esters (PAEs). They are globally produced in huge quantities each year for a wide range of applications being the main plasticizers used in the polymer industry since the 1930s (Liang et al., 2008). They are commonly added (10% - 60%) to plastic materials, such as polyvinyl chloride (PVC), polyethylene terephthalate (PET), polyvinyl acetate (PVA), and polyethylene (PE), to improve extensibility, elasticity, and workability of the polymers. Phthalates are considered as priority industrial pollutants due to their chemical toxicity, and adverse effect on human health and animals (Giuliani et al., 2020). In environment, degradation of phthalate esters by bacteria involves the initial de-esterification step which releases readily degradable side chain alcohols and often accumulates phthalate. Decarboxylation of phthalate is a challenging and rate-limiting reaction in microbial degradation, especially for anaerobic bacteria (Kleerebezem et al., 1999). Because, aerobic phthalate decarboxylation is facilitated by oxygenase-dependent oxygenation reactions, forming a 3,4-dihydrodiol (in Gram-positive bacteria) or 4,5-dihydrodiol (in Gram-negative bacteria) which is later converted to dihydroxyphthalate which is then decarboxylated by decarboxylases to the common intermediate 3,4-dihydroxybenzoate (Batie et al., 1987; Chang & Zylstra, 1998; Eaton & Ribbons, 1982). Similarly, *m*-phthalate (isophthalate) and *p*-phthalate (terephthalate) are analogously decarboxylated leading to 3,4-dihydroxybenzoate (Fukuhara et al., 2010; FUKUHARA et al., 2008; Karegoudar & Pujar, 1985; Schläfli et al., 1994). In essence, aerobic phthalate-degrading bacteria introduce molecular oxygen into the phthalate ring that partially polarizes the ring, facilitating the difficult step of phthalate decarboxylation.

Due to absence of molecular oxygen, decarboxylation in anaerobic phthalate-degrading bacteria becomes challenging and regarded as the rate-limiting step (Kleerebezem et al., 1999). In the past, different hypotheses were proposed for anaerobic decarboxylation of phthalate to benzoate. For instance, Taylor and Ribbon (1983) suggested that phthalic acid is reduced by two electrons leading to 3,5-cyclohexadiene-1,2-dicarboxylic acid before its decarboxylation to benzoic acid (Taylor & Ribbons, 1983). Later, it was assumed that decarboxylation of phthalate involves formation of phthalate-coenzyme A ester, which is subsequently decarboxylated to benzoyl-CoA (Nozawa & Maruyama, 1988a; Nozawa & Maruyama, 1988b). In recent years, we provided the first experimental evidence that phthalate degrading anaerobic bacteria accomplish phthalate decarboxylation via a two-step enzyme reaction involves activation of *o*-phthalate to *o*-phthalyl-CoA, which is subsequently decarboxylated to benzoyl-CoA (Boll et al., 2002; Geiger et al., 2019; Junghare et al., 2019; Junghare et al., 2016). We demonstrated the formation of first intermediate *o*-phthalyl-CoA from *o*-phthalate and succinyl-CoA followed its decarboxylation to benzoyl-CoA in assay performed in cell-free extract of *Azoarcus* sp. strain PA01 (Junghare et al., 2016). Benzoyl-CoA is further metabolized by the enzymes of the anaerobic benzoyl-CoA degradation pathway (Breese et al., 1998; Fuchs, 2008; Gall et al., 2013). The genome of *Azoarcus* sp. strain PA01 possess the required genes for anaerobic benzoate-degradation pathway (Junghare et al., 2015).

Differential protein profiling using cell-extracts of benzoate-grown cells versus o-phthalate of *Azoarcus sp.* strain PA01 identified a set of proteins induced specifically with o-phthalate. Interestingly, phthalate-induced proteins were placed together in a single gene cluster that includes genes coding for the protein homologous to a solute transporter (locus tag PA01_00214), two CoA-transferases (PA01_00215 and PA01_00216), and the UbiD-like/UbiX-like decarboxylases (PA01_00217 and PA01_00218), respectively (Junghare et al., 2016). In phthalate degrading denitrifying bacteria, the CoA-transferases are involved in *o*-phthalate activation to *o*-phthalyl-CoA, and UbiD-like/UbiX-like decarboxylases in decarboxylating *o*-phthalyl-CoA to benzoyl-CoA (Ebenau-Jehle et al., 2017; Junghare et al., 2016; Mergelsberg et al., 2018). Among these, the genes involved in activation of phthalate to phthalyl-CoA were belonged to CoA transferase or CoA ligase family that were homologous to proteins involved in activation of aromatic acids to its CoA esters.

Interestingly, in all phthalate degrading bacteria including *Azoarcus* sp. PA01 the genes involved in decarboxylation were annotated as proteins homologous to UbiD-like/UbiX-like decarboxylases. UbiD-like and UbiX-like are proteins that widely distributed among prokaryotes, e.g., *Escherichia coli* with the carboxy-lyase activity converting 4-hydroxy-3-octaprenyl benzoic acid to 2-octaprenylphenol, an early step in ubiquinone biosynthesis (Gulmezian et al., 2007; Lin et al., 2015; Meganathan, 2001; White et al., 2015). Even though 4-hydroxy-3-octaprenyl benzoic acid and *o*-phthalyl-CoA are structurally quite different from each other, the genes PA01_00217 and PA01_00218 that are homologous to UbiD-like/UbiX-like proteins are suggested to be involved in the anaerobic decarboxylation of phthalyl-CoA to benzoyl-CoA by *Azoarcus* sp. strain PA01 (Junghare et al., 2016). To address the specific role of UbiD-like and UbiX-like decarboxylase, especially the question how these two proteins function together to catalyse the decarboxylation of *o*-phthalyl-CoA to benzoyl-CoA. We performed the cloning, expression and characterization of both the proteins UbiD-like/PhtDa (60 kDa) and UbiX-like/PhtDb (23 kDa). In our preliminary results we could provide the first biochemical evidence that both the proteins are involved in the decarboxylation of *o*-phthalyl-CoA to benzoyl-CoA, of which UbiX-like protein prepares modified prFMN co-cofactor that is required by UbiD-like for actual decarboxylation during the anaerobic degradation of *o*-phthalate by *Azoarcus* sp. strain PA01.

## Results

### Gene amplifications, cloning of PhtDa and PhtDb

Genomic DNA used for amplification of full-length *o*-phthalyl-CoA decarboxylase genes resulted in a 1.58 kb amplicon for PA01_00217 and 0.6 kb amplicon for PA01_00218 (Figure 1A). PCR products were cloned into the ampicillin resistance pET100/D-TOPO expression vector which adds 6His-tag (3kDa), Xpress Epitope to the N-terminal end of the overexpressed protein. Recombinant plasmid was used to transform *E. coli* TOP10 cells and grown on LB media containing ampicillin (100 µg/ml). Several positive clones were found with a gene of interest inserted into the pET100/D-TOPO expression vector. Recombinant clones were picked and screened based on the PCR product size of amplified PCR products using T7 primers (as discussed in method section) and confirmed by comparing the size of the respective genes plus additional 276 bp from cloning site of pET100/D-TOPO vector (Figure 1AB). The gene coding for PhtDa is 1584 bp long plus 276 and PhtDb is 600 bp plus 276 and expected size PCR products (using T7 primers) from positive clones are shown in Figure 1B.

**Figure 1.**
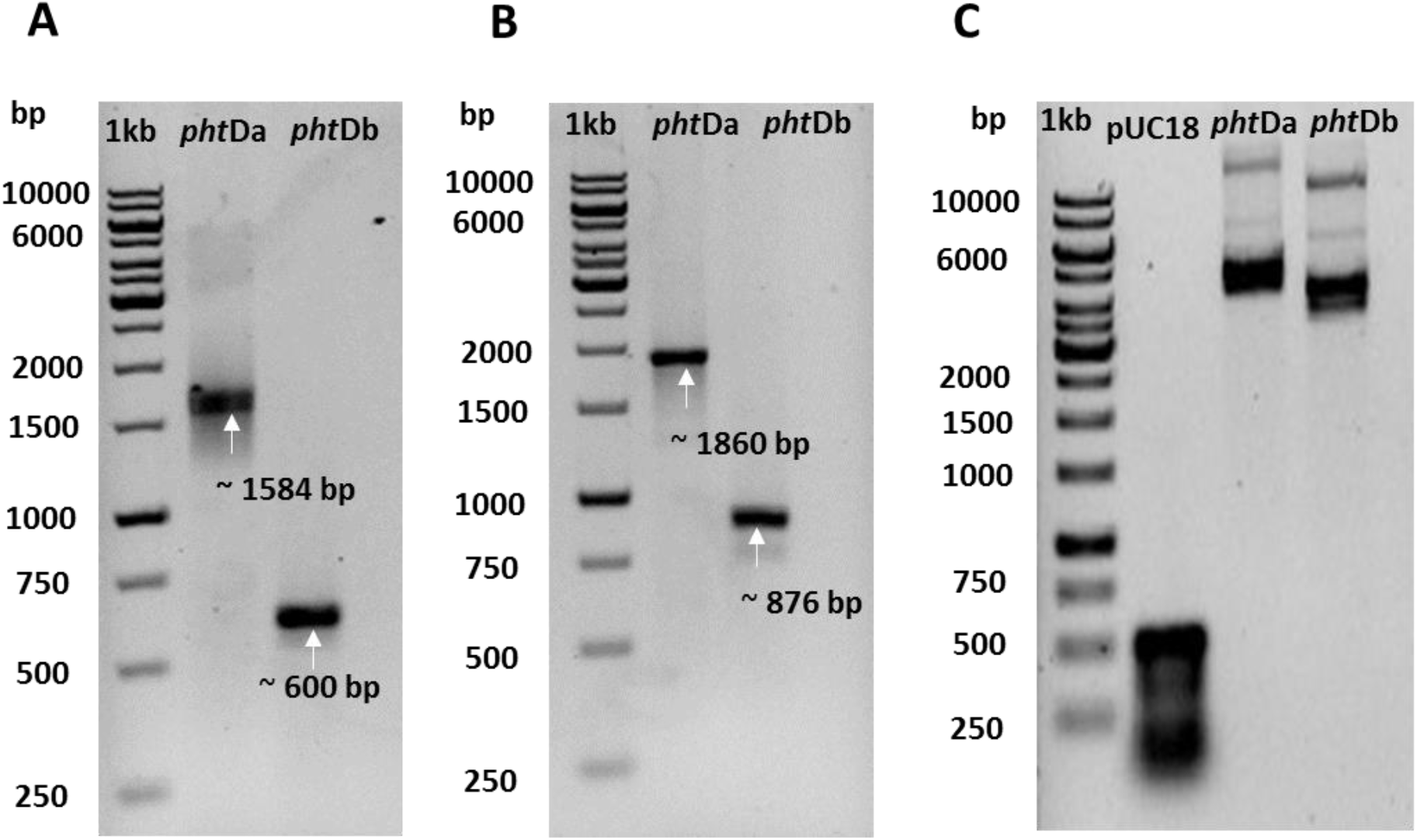
Electrophoretic analysis of PCR-amplified genes in 1.0 % agarose gel stained with ethidium bromide: **A)** PCR amplification of genes PA01_00217, 1.58 kb and PA01_00218, 0.6 kb from gDNA of *Azoarcus* sp. strain PA01; **B)** screening of the positive clones for insertion of the respective genes into the pET100/D-TOPO expression vector PCR amplified using T7 primers (expected size of the PCR amplicon is calculated from original gene length plus 276 bp from cloning site of vector); and **C)** recombinant plasmid DNA isolation. 1 kb gene ruler (Thermo Fischer) was used to determine the molecular size of the PCR products on the gel.

Selected positive clones of *E. coli* TOP10 cells containing recombinant plasmids were propagated in LB media (ampicillin, 100 μg/ml) for plasmid DNA isolation. Purified recombinant plasmid DNAs were sequenced across the cloning site using T7 primers and inserted gene sequences were determined (data not shown). The sequence of recombinant genes displayed correct insertion of genes into the vector and showed 100% sequence identity with the original gene sequences available from the published draft genome sequence of *Azoarcus* sp. strain PA01 (Junghare *et al*., 2015b).

### Heterologous expression, purification and protein identification

The genes PA01_00217 and PA01_00218 (Figure 1A), putatively coding for the enzymes PhtDa and PhtDb responsible decarboxylation of *o*-phthalyl-CoA to benzoyl-CoA, i.e., *o*-phthalyl-CoA decarboxylase, were individually expressed in *E. coli* BL21 star (DE3) and *E. coli* Rosetta 2 (DE3)pLysS host cells with a N-terminal His-tag introduced from pET expression vector. The purified recombinant pET100/D-TOPO plasmids (Figure 1C) containing genes coding for PhtDa (60 kDa) and PhtDb (60 kDa) were used to transform the chemically competent *E. coli* BL21 star (DE3) and *E. coli* Rosetta 2 (DE3)pLysS as the expression host cells and grown on LB media with ampicillin selection (100 µg/ml). Complete open reading frames of the cloned genes PA01_00217 and PA01_00218 are expected to encode the proteins of 527 (PhtDa) and 199 (PhtDb) amino acids in length with the computed molecular weight of about 59.54 kDa (̴60 kDa) and 21.99 kDa (̴22 kDa), respectively, plus 3 kDa 6His-tag.

In order to find the optimal expression conditions for obtaining soluble proteins, both the transformed *E. coli* expression host strains were grown under different combinations of growth conditions, including temperatures (37, 25 and 15 °C), culture media (LB and TB medium) varying IPTG concentrations (0.1 to 1 mM) and with or without glucose (1 %) plus ethanol (3 %), respectively, (data not shown). Protein expression optimization experiments revealed, *E. coli* Rosetta 2 (DE3)pLysS host cells were found to be suitable for optimal expression of soluble protein after induction with 0.5 mM IPTG (at OD between 0.5 - 0.6) and grown at 15 °C for about 18 - 20 h in the TB medium supplemented with 3 % ethanol and 1 % glucose (Figure 2A and 2B). Although, under these growth conditions protein over-expression was also observed at 37 °C and 25 °C, but expressed proteins formed inclusion bodies (insoluble protein). Finally, *E. coli* Rosetta 2 (DE3)pLysS cells cultured in TB media was selected for the large scale protein expression and purifications. Protein expression was up-scaled to 500 - 1000 ml culture volume.

**Figure 2.**
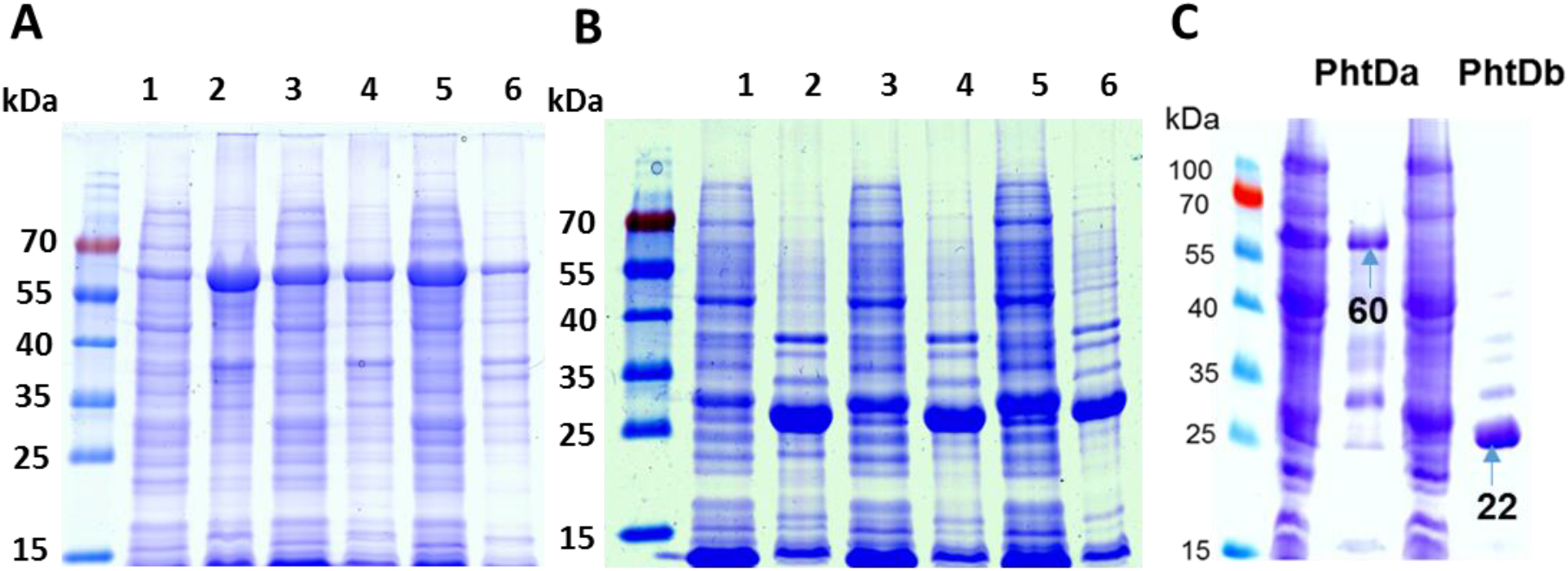
Sodium dodecyl sulfate-polyacrylamide (12 % resolving) gel electrophoretic analysis of proteins expressed in *E. coli* Rosetta 2 (DE3)pLysS cells grown in different culture media (3 % ethanol and 1 % glucose) and induction with 0.5 mM of IPTG at varying temperatures: **A)** expression level of PhtDa protein (̴60 kDa); and **B)** expression level of PhtDb protein (̴ 23 kDa): lane 1 - 2, cell-extract and pellet of cells grown at 37 °C for 4 h; lane 3 - 4, cell-extract and pellet of cells grown at 25 °C for 7.5 h; lane 5 - 6, cell-extract and pellet of cells grown at 15 °C for 20 h; and **C)** partial purification of PhtDb (̴ 60 kDa ± 3) and PhtDb (̴ 22 kDa ± 3) proteins using the Ni^2+^ ion affinity column. The pre-stained protein molecular size marker (range 15 – 100 kDa) was used to estimate the size of the protein bands.

His-tag on the recombinant proteins (PhtDa and PhtDb) allowed quick and easy purification from the supernatant. About 6 - 8 ml of supernatant prepared from 3 - 4 g of wet cell material lysate was used to load the Ni^2+^ ion chelated column (40 mg His-tagged protein binding capacity). By washing the Ni^2+^ ion chelated column with about 10 ml buffer A (as discussed above), his-tagged proteins were eluted with buffer B containing higher concentrations of imidazole (0.5 M). Eluted fractions were subjected to SDS-PAGE analysis and simultaneously, purified protein contents were determined using the Bradford assay. SDS-PAGE analysis of Ni^2+^ ion affinity chromatography purified proteins resulted the dense single protein bands at the expected size of about 62 kDa for PhtDa and 23 kDa for PhtDb that include plus 3 kDa 6His-tag (Figure 2C). Protein bands excised from the SDS gel and identified peptides by MS analysis, showed the expected molecular mass and high identity with the deduced amino acid sequence of genes PA01_00217 and PA01_00218 from the draft genome of the *Azoarcus* sp. strain PA01 (data not shown).

### Enzyme activity, substrate specificity and cofactor requirement

Decarboxylase activity for conversion of *o*-phthalyl-CoA to benzoyl-CoA was determined with recombinantly expressed PhtDa and PhtDb in *E. coli*. Enzyme assay were performed directly with the *E. coli* transformant’s cell lysate or cell-free extracts containing respective expressed proteins PhtDA and PhtDb. Decarboxylation of *o*-phthalyl-CoA was demonstrated with *E. coli* cell-free extracts expressing PhtDa and PthDb proteins and LC-MS analysis of samples collected at the different time intervals revealed an increase of benzoyl-CoA (Figure 3). Decarboxylase activity was unaffected by oxygen exposure and was independent of oxygen. Neither one of these proteins could decarboxylate *o*-phthalyl-CoA to benzoyl-CoA alone. As expected, no decarboxylase activity was observed with isophthalyl-CoA or terephthalyl-CoA (Junghare *et al*., 2016). Furthermore, decarboxylase activity was observed only when, PhtDb was previously incubated with flavin mononucleotide (FMN) and dimethylallyl monophosphate (DMAP). No activity was observed when FMN was replaced with the flavin dinucleotide (FAD). This result strongly indicates that PhtDb is a FMN-binding/dependent protein that is essential for the decarboxylation of *o*-phthalyl-CoA to benzoyl-CoA. In addition to this, UV-visible scanned spectrum of FMN-DMAP incubated and His-tag purified PhtDb revealed the possible binding and modification of the FMN molecule to a PhtDb protein (Figure 4). UV-visible scanning of PhtDa and PhtDb showed the same absorption maxima at the wavelength of about 340 nm but at different intensity.

**Figure 3.**
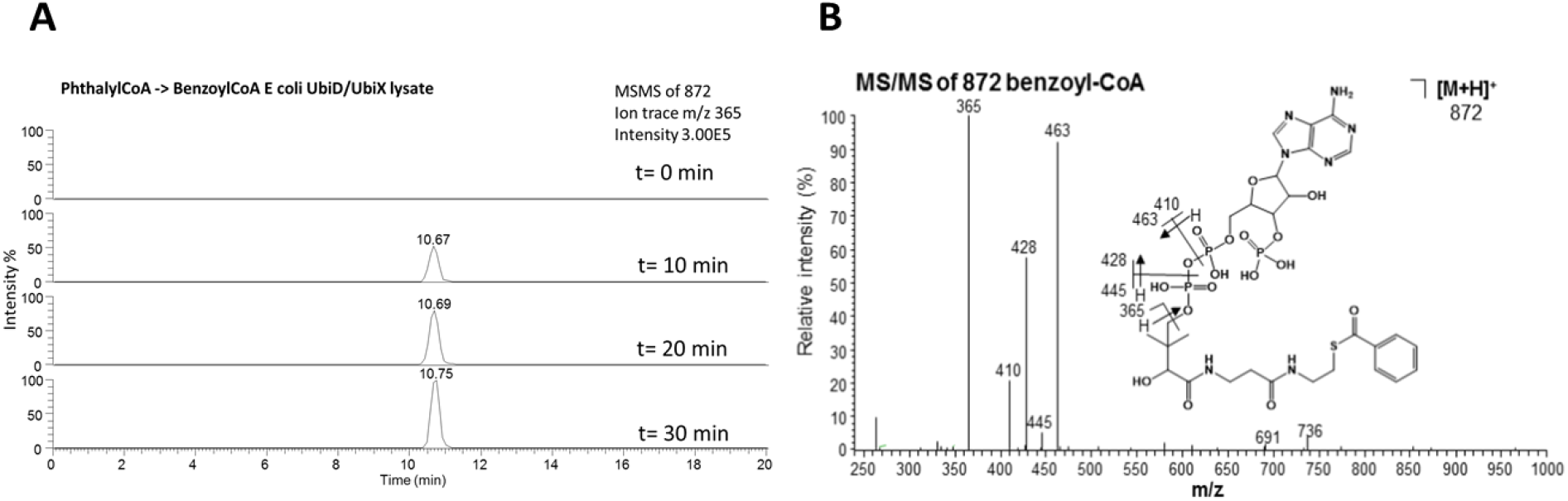
LC-MS/MS ion traces following benzoyl-CoA formation from the *o*-phthalyl-CoA decarboxylation in an enzyme assay performed with cell-free extracts of recombinantly expressed PhtDa and PhtDb in *E. coli*. **A)** Time course of formation of benzoyl-CoA from *o*-phthalyl-CoA monitoring the specific ion traces m/z 365 of the MS/MS fragmentation of benzoyl-CoA (quasimolecular ion m/z 872); and **B)** ESI-MS/MS of the quasimolecular ion m/z 872 of benzoyl-CoA.

**Figure 4.**
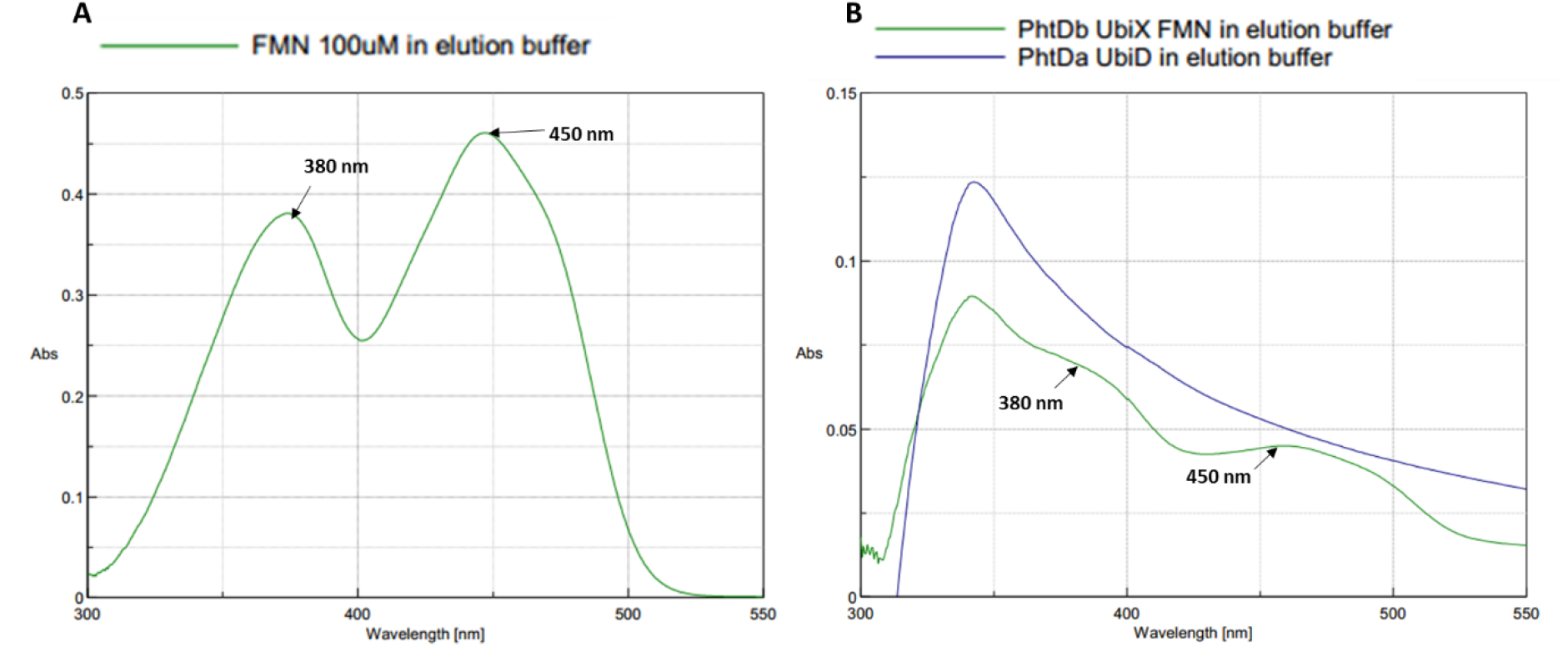
UV-visible scanned spectrum of: a) flavin mono nucleotide (FMN) in elution buffer; b) FMN-DMAP incubated and His-tag purified PhtDb showing absorption peak maxima at a wavelength of about 340 nm, 380 nm and 458 nm possible indicating binding of FMN molecules (preliminary results). PhtDa without FMN was scanned as the control.

### Phylogenetic position of PhtDa and PhtDb

A phylogenetic tree was constructed using MEGA 7.0 software (Kumar *et al*., 2016) based on the neighbor-joining method to determine the evolutionary relationship of PhtDa and PhtDb of *Azoarcus* sp. strain PA01 with the other known bacterial and fungal species UbiX-like/UbiD-like decarboxylases whose crystal structure are available. Amino acid sequence of total 13 different proteins were aligned with clustal W method and a single phylogenetic tree was generated for the evolutionary analysis (Figure 5). Among all the UbiD-like and UbiX-like enzymes from the bacterial and fungal species, the PhtDa enzyme (PA01_00217) with expected decarboxylase activity formed a discrete group with UbiD-like and a putatively flavin (FMN)-binding PhtDb (PA01_00218) grouped with UbiD-like enzymes, respectively (Figure 5). The proteins PhtDa and PhtDb were clearly separated from each other in the distinct clade and showed only about 30 % sequence identity to one another. Among the UbiD-like enzymes clade in the phylogenetic tree, PhtDa of *Azoarcus* sp. strain PA01 was found to be more closely related with the putative aromatic acid decarboxylase (Pad1, 4IP2_C) of *P. aeruginosa* and the ferulic acid decarboxylase (Fdc1, AHY75481) of *S. cerevisiae* showing about 23 % of sequence similarity. Whereas the PhtDb FMN-binding protein of *Azoarcus* sp. strain PA01 shared highest 64 % of amino acid sequence similarity with UbiX-like enzyme (P69772) of *E. coli* strain O157:H7 (Figure 5). The other closely related UbiX-like enzymes includes the phenolic acid decarboxylase (β-subunit, P94404) of *B. subtilis*, fungal phenylacrylic acid decarboxylases (Pad1/PadA1) of *S. cerevisiae* (P33751) and *A. niger* (A3F715) were positioned relatively distant with around 50 % amino acid sequence similarity, but all grouped together UbiX-like enzymes (Figure 5). However, the UbiX (Q9HX08) flavin prenyltransferase enzyme of *P. aeruginosa* strain PAO1 showed only about 44 % sequence identity with PhtDb and was distantly placed compared other UbiX (Figure 5).

**Figure 5.**
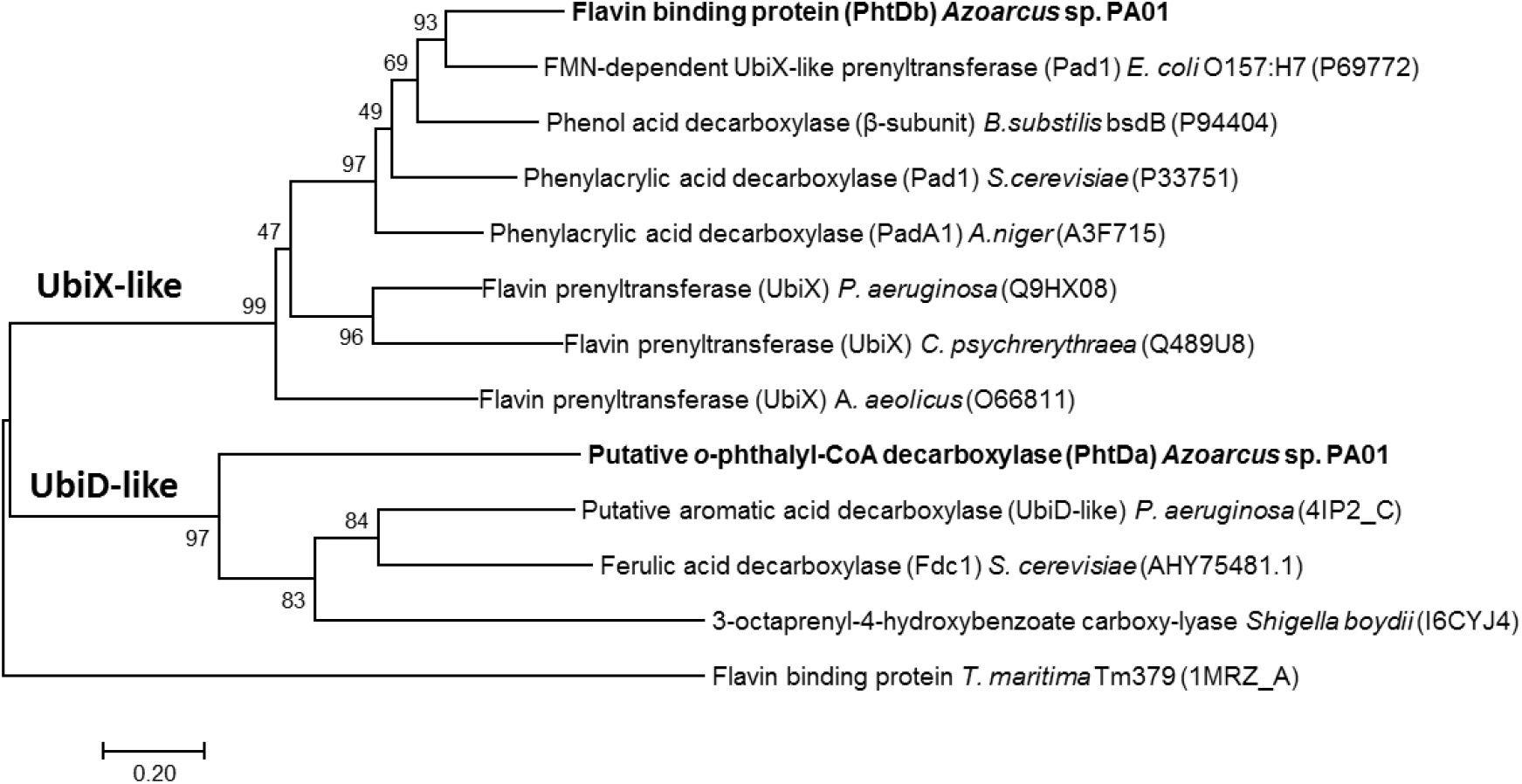
Phylogenetic tree showing the relationship of PhtDa and PhtDb proteins of *Azoarcus* sp. strain PA01 that are involved in the decarboxylation of *o*-phthalyl-CoA to benzoyl-CoA and the other known UbiX-like and UbiD-like enzyme from bacterial and fungal species. Bar indicates 2 % sequence divergence.

### Multiple sequence alignment and homology-based structure modelling

A BLAST search for proteins PhtDa and PhtDb found several sequences sharing significant similarity (data not shown). However, the BLAST results contain the enzymes with known functions that shared less similarity with PhtDa and PhtDb. For example, the phylogenetic analysis of PhtDa suggested, it shared only about 23 % sequence identity with the UbiD-like decarboxylase (Pad1) of *P. aerugonisa* (PDB code 4IP2_C, UniProt Q9I6N5) whose crystal structure is currently known (Jacewicz *et al*., 2013). Therefore, due to very less sequence similarity (23 %) of PhtDa to enzyme with known function, multiple sequence alignment and secondary structure prediction for PhtDa was not performed. On the other hand, BLAST search and phylogenetic analysis of PhtDb revealed about 64% sequence identity with FMN-dependent UbiX-like decarboxylase of *E. coli* O157:H7 (UniProt P69772) whose crystal structure (PDB code 1sbz) is currently resolved (Rangarajan *et al*., 2004). Other known UbiX-like enzymes of bacterial and fungal species also shared significant identity with PhtDb (Figure 5). Therefore, multiple amino acid sequence alignment and secondary structure predictions (using PDB code 1sbz as template) of PhtDb revealed highly conserved regions not only with the FMN-dependent UbiX-like decarboxylase of *E. coli* (P69772), but also with other UbiX-like enzymes from bacterial and fungal species (Figure 5). We have analyzed more closely the top 5 sequences, which all had lengths of ∼200 residues and had over 50 % sequence identity to PhtDb of *Azoarcus* sp. strain PA01. Among the ∼200 amino acid residues in the sequences, there are ∼50 residues conserved in all sequences, as highlighted (red) as shown in Figure 6. The majority of these enzymes sequences are the known FMN-binding prenyltransferase belonging to the superfamily of UbiX-like enzymes.

**Figure 6.**
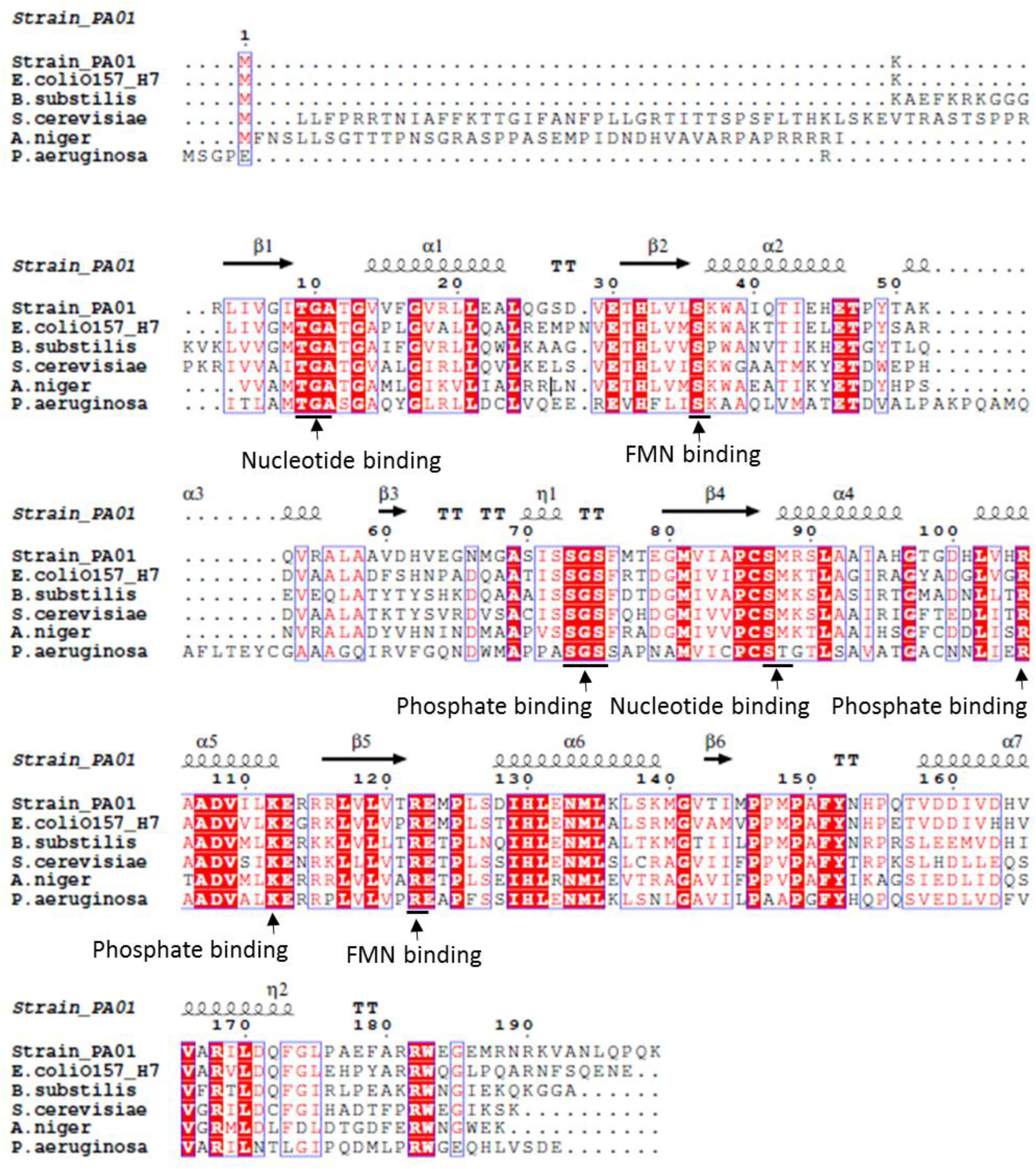
Multiple sequence alignments of PhtDb (PA01_00218) of *Azoarcus* sp. strain PA01 with other UbiX-like enzymes from selected bacterial or fungal species: *E. coli* O157:H7 (P69772), *B. subtilis* bsdB (P94404), *S. cerevisiae* Pad1 (P33751), *A. niger* PadA1 (A3F715) and *P. aeruginosa* UbiX (Q9HX08). Conserved amino acid residues that putatively involved in flavin mononucleotide nucleotide (FMN), phosphate, and nucleotide binding are indicated by labelled arrows. The predicted secondary structure elements of PhtDb are shown based on the ESPrip 3.0 online tool. Alpha (α) helices and 3_10_-helices (denoted as ղ) are shown as squiggles, β-strands as arrows and β-turns as TT. Conserved amino acid sequences of PhtDb are presented as highlighted regions and red denotes highly conserved amino acids regions.

The three-dimensional (3-D) structures of many UbiD-like and UbiX-like enzymes are presently known. The 3-D structure prediction and modelling of PthDa and PhtDb were computed in SWISSMODEL workspace (https://swissmodel.expasy.org/interactive) using the deduced amino acid sequences of the PhtDa (PA01_00217) and PhtDb (PA01_00218) to find the best templates. Although, amino acid sequence of ferulic acid decarboxylase (Fdc1) from *S. cerevisiae* and putative aromatic acid decarboxylase (Pad1) of *P. aeruginosa* shared ∼23% amino acid sequence similarity with PhtDa (Figure 5). The homodimer crystal structure of Pad1 (PDB code 4iws) of *P. aeruginosa* (Jacewicz *et al*., 2013) was the best template and chosen for homology or template based structural modelling of PhtDa (Figure 7A). Whereas the homo 12-mer (dodecamer) crystal structure (PDB code 1sbz) of FMN-binding UbiX flavoprotein (Pad1) in *E. coli* O157: H7 (Rangarajan *et al*., 2004) was the best template for PhtDb and chosen for predicting the 3-D structure of PhtDb (Figure 7B). The predicted 3-D structures of PhtDa (homodimer) and PhtDb (homo 12-mer) in *Azoarcus* sp. strain PA01 were visualised (Figure 7AB) and compared and superimposed with the respective templates (PDB code 4zac and 1sbz) using the UCSF-Chimera package (Figure 7CD). PhtDb showed a high degree of secondary structure and motif conservation with the template (Figure 7D), compared to overlapping of PhtDa with its template (Figure 7C).

**Figure 7.**
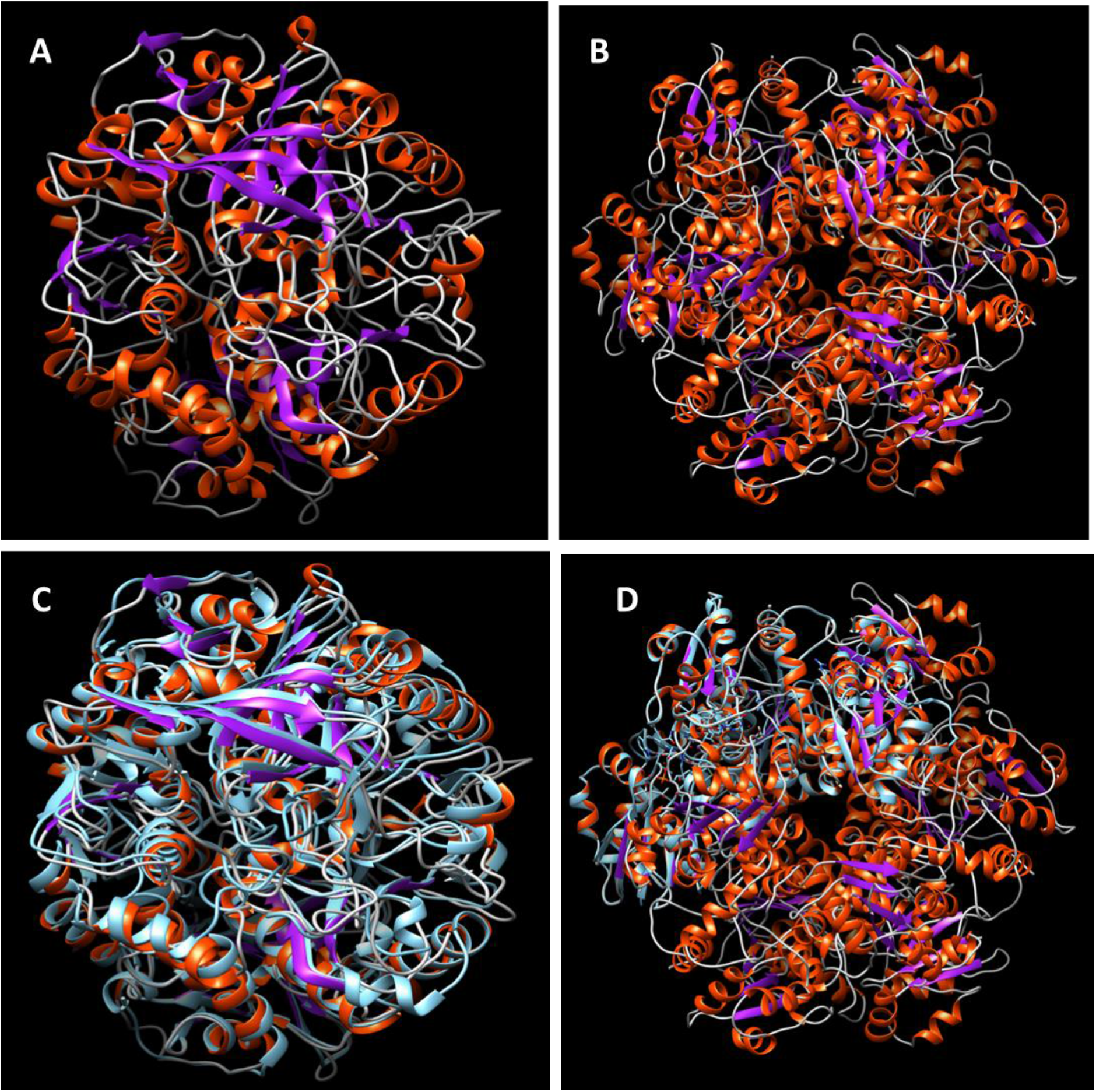
Homology-based molecular modelling of PhtDa (PA01_00217) and PhtDb (PA01_00218) proteins with the decarboxylase activity from *Azoarcus* sp. strain PA01 were constructed using SWISSMODEL workspace: **A)** a 3-D model (homodimer) structure of PhtDa based on the homodimer crystal structure (PDB code 4iws) of Pad1 in *P. aeruginosa* (Jacewicz *et al*., 2013); **B)** a 3-D model (homo 12-mer) of PhtDb based on the crystal structure (PDB code 1sbz) of UbiX flavoprotein (Pad1, homo 12-mer) in *E. coli* O157:H7 as a template (Rangarajan *et al*., 2004). Superimposition of predicted 3-D model structure of: **C)** PhtDa, and **D)** PhtDb with their respective templates (4iws and 1sbz) are shown in faint blue color performed using UCSF-Chimera package (https://www.cgl.ucsf.edu/chimera/).

### Native PAGE and protein identification by MS analysis

In order to investigate the protein complexes in phthalate-induced proteins by *Azoarcus* sp. strain PA01, especially to determine the PhtDa and PthDb protein-protein interactions and oligomer state. Native-protein gel electrophoresis was performed with cell-free extracts of *o*-phthalate *versus* benzoate-grown cells. Interestingly, only few native protein bands were visible on *o*-phthalate-grown cells compared to benzoate-grown cells (Figure 8A). Specifically phthalate-induced protein bands were estimated between 400 to 140 kDa that include protein bands labeled as 2, 3, 4 and 5 (Figure 8A). These spots were excised from the native gel and identified by MS. Protein identification of protein bands 2 and 3 (approximate estimated molecular size ranging between 390 to 350 kDa) resulted the identification majorly two proteins, namely PhtDa (60 kDa) and PhthDb (22 kDa) as shown in Table 1. Among the protein identification of the lower molecular weight protein bands (4, 5, 6 and 7) PhtDa was also identified, including the CoA-transferases (PhtSa and PhtSb protein) that are involved in the activation of *o*-phthalate to *o*-phthalyl-CoA reported in our recent findings (Junghare *et al*., 2016).

**Figure 8.**
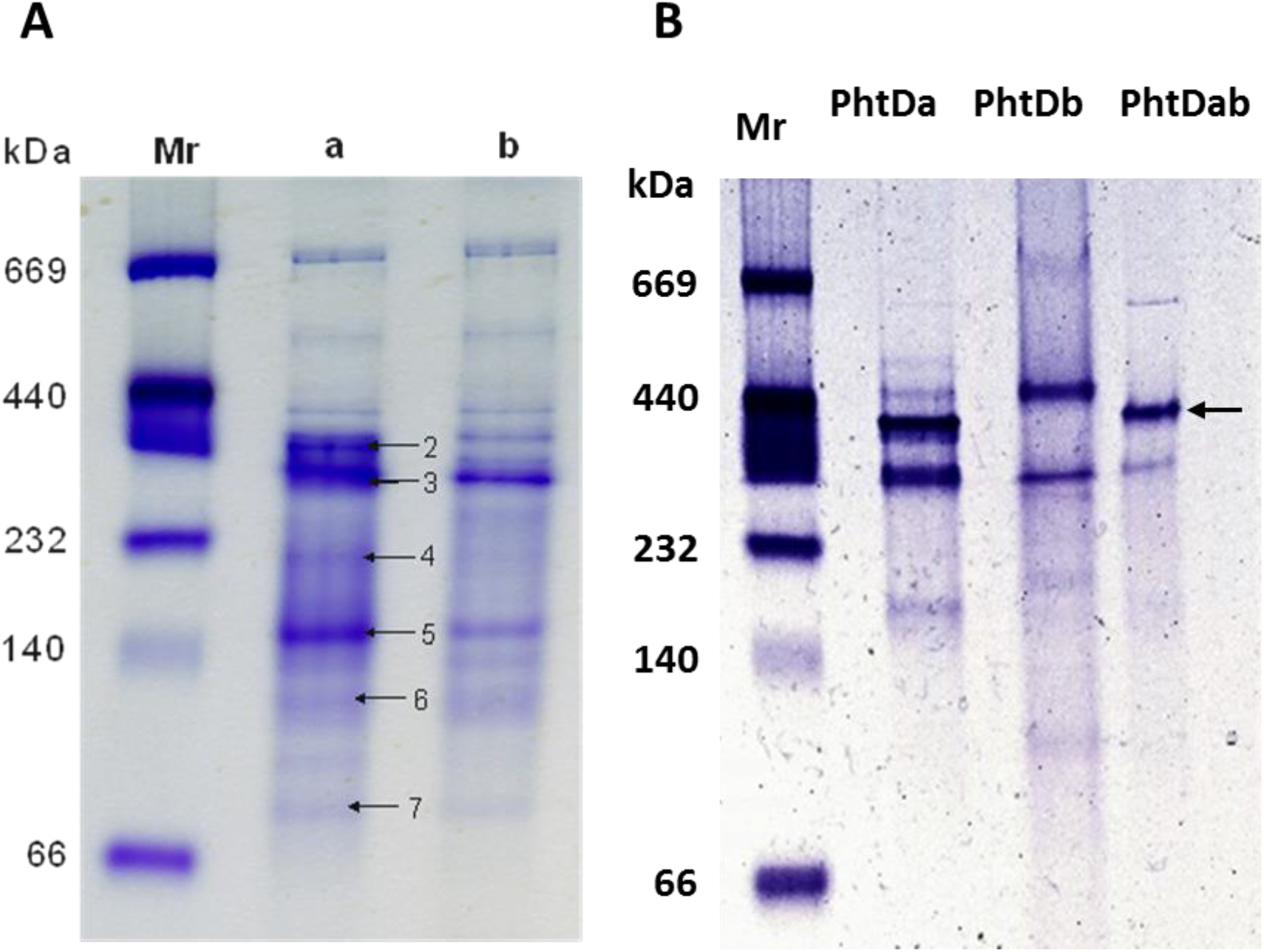
Native-PAGE analysis of: **A)** the soluble protein fraction from cell-free extracts of *Azoarcus* sp. strain PA01 cells grown with a) o-phthalate and b) benzoate; and **B)** partially purified PhtDa, PhtDb and PhtDab expressed in *E. coli*. Protein separation was performed using the Mini-PROTEAN TGX native gel slab (BioRad) and bands were visualized by staining with Coomassie Brilliant Blue G250. Lane Mr: native high molecular weight protein marker (Amersham, GE Healthcare life sciences). The arrow in figure B highlights the probable protein complex of PhtDa and PhtDb of estimated size of about 390 kDa (PhtDa 60 kDa x dimer = 120 kDa plus PhtDb 22 kDa x 12 mer = 264 kDa).

**Table 1.** Protein identification by mass spectrometry of the spots excised from native PAGE from the *o*-phthalate-grown cells compared to benzoate grown cells of *Azoarcus sp.* strain PA01.

| Band No. | MW on gel (kDa) | Locus tag | Major proteins identified by MS analysis | Calculated MW (kDa) | Coverage % | Score |
| --- | --- | --- | --- | --- | --- | --- |
| 2 | 390 | 00217 | PhtDa, UbiD-like | 58.8 | 74.57 | 15680 |
|  |  | 00218 | PhtDb, UbiX-like | 22.0 | 53.27 | 616 |
| 3 | 350 | 00217 | PhtDa, UbiD-like | 58.8 | 70.02 | 8169 |
|  |  | 00218 | PhtDb, UbiX-like | 22.0 | 56.28 | 418 |
| 4 | 210 | 00217 | PhtDa, UbiD-like | 58.8 | 59.96 | 5812 |
| 5 | 140 | 00217 | PhtDa, UbiD-like | 58.8 | 61.29 | 10500 |
|  |  | 00216 | PhtSb, CoA-transferase | 43.6 | 14.74 | 194 |
|  |  | 00215 | PhtSa, CoA-transferase | 43.8 | 14.04 | 114 |
| 6 | 100 | 00217 | PhtDa, UbiD-like | 58.8 | 49.72 | 2101 |
|  |  | 00215 | PhtSa, CoA-transferase | 43.8 | 20.20 | 261 |
|  |  | 00216 | PhtSb CoA-transferase | 41.3 | 13.65 | 88.85 |
| 7 | 80 | 00215 | PhtSa, CoA-transferase | 43.8 | 53.69 | 2019 |
|  |  | 00216 | PhtSb CoA-transferase | 41.3 | 47.77 | 1096 |
|  |  | 00217 | PhtDa, UbiD-like | 58.8 | 33.97 | 661 |

Note- From the each spot different proteins were identified. However, in this table the results of four proteins (four subunits) that are involved in the initial steps of *o*-phthalate degradation were shown. \*SPA stands for spots from *o*-phthalate-grown gel, SBA stands for spots from benzoate-grown gel and MMPA are spots from membrane proteins of *o*-phthalate-grown *Azoarcus* sp. PA01.

Furthermore, native-protein gel analysis of heterologously expressed and partially purified PhtDa and PhtDb, were performed individually and incubating together for about 1 h at room temperature to determine the protein-protein interactions. Surprisingly, when PhtDa (60 kDa) and PhtDb (22 kDa) were incubated together with FMN and DMAP, only a single band at ca. 400 kDa was visible on the native protein gel (Figure 8B). Furthermore, individual native protein gel analysis of purified PhtDa and PhtDb revealed the formation of two distinct bands visible at the estimated molecular size between ca. 280 to 420 kDa. Nevertheless, SDS analysis of purified PhtDa (60 kDa) and PhtDb (22 kDa) showed only a single bands at the expected size for the respective proteins (Figure 2C). These results strongly indicate that these high molecular size protein bands in native gel are most likely the multimeric complexes of PhtDa and PhtDb. However, at the moment it is still unclear whether the single band observed in the lane of the PhtDab incubated mixture (with FMN plus DMAP) is due to the PhtDa and PhtDb multimeric complex, because MS confirmation of the proteins is elusive.

## Discussion

Present work demonstrates that PhtDa is the enzyme responsible for catalysing decarboxylation of *o*-phthalyl-CoA to benzoyl-CoA, and the decarboxylase activity of PhtDa depends on the FMN-dependent PhtDb protein constituting a two-enzyme component system. PhtDa (60 kDa) and PhtDb (22 kDa) were cloned into pET100/D-TOPO and overexpressed as N-terminal His-tagged proteins in *E. coli* Rosetta (DE3)pLysS host. Both proteins were purified to homogeneity by utilizing Ni^2+^ ion affinity chromatography (Figure 2C). The molecular weight of the recombinant proteins was determined by SDS PAGE analysis. Alternatively, proteins were identified by MS analysis and revealved that both protein bands showed expected molecular masses and high identity with the deduced amino acid sequenes of both gene from the genome sequence of *Azoarcus* sp. strain PA01 (Junghare *et al*., 2015b).

Enzyme assays with cell lysate or cell-free extracts of *E. coli* strains expressing PhtDa and PhtDb could readily convert *o*-phthalyl-CoA to benzoyl-CoA (Figure 3). Interestingly, none of these proteins possess decarboxylase activity alone. Although, PhtDa and PhtDb do not share sequence similarity with each other, but they belong to a new class of enzyme family and are homologous to UbiD-like and UbiX-like decarboxylases found in a wide range of bacterial and fungal species (Figure 5, 6 and 7). UbiD and UbiX are known to act in the decarboxylation of aromatic substrates (Zhan and Javor, 2003) but individually UbiX or UbiD were unable to perform decarboxylation (Jacewicz *et al*., 2013; Rangarajan *et al*., 2004; Kopec *et al*., 2011). Our results showing the involvement of PhtDa and PhtDb for decarboxylation reaction are in agreement with the previous studies. Because decarboxylation of phenylacrylic acid in *S. cerevisae* required Fdc1 (UbiD-like) and Pad1 (FMN dependent UbiX-like). Similarly, in *E. coli* O157:H7 or *P. aeruginosa* decarboxylation of 3-octaprenyl-4-hydroxybenzoate to 2-octaprenylphenol is also required two enzymes as UbiX-like (FMN-dependent) and an UbiD-like aromatic acid decarboxylase (Rangarajan *et al*., 2004; Jacewicz *et al*., 2013).

Furthermore, heterologously overexpressed PhtDa and PhtDb in *E. coli* only led to benzoyl-CoA formation, when PhtDb was pre-incubated with FMN and dimethylallylphosphate (DMAP). No decarboxylase activity was observed when FMN was replaced against flavin dinucleotide (FAD). These results strongly suggest that decarboxylase activity of PhtDa, is dependent on FMN-dependent PhtDb. Phylogenetically, PhtDb is closely related to the FMN-dependent UbiX-like enzyme which covalently modifies FMN molecule that is needed for the decarboxylase activity by UbiD-like decarboxylase in *E. coli* (White *et al*., 2015; Lin *et al*., 2015; Rangrajan *et al*., 2015; Payne *et al*., 2015). Multiple sequence alignment of PhtDb with the known UbiX flavin-binding proteins from bacterial and fungal species (Figure 5 and 6) revealed highly conserved amino acid residues. PhtDb showed highest sequence similarity (64 %) with the FMN-dependent UbiX-like enzyme of *E. coli* O157:H7 for which the FMN-binding site at the amino acid positions Ser36 and Arg122, and a nucleotide binding site at Thr9, Gly10, Ala11, Ser87 and Met88, respectively, has been established recently (White *et al*., 2015). Interestingly, PhtDb also possesses the conserved amino acid residues at these positions, suggesting a probable binding site for FMN and DMAP along with the similar FMN dependent enzymes (Figure 6). The addition of DMAP and FMN to PhtDb caused distinct changes in the FMN absorbance profile (Figure 4), which indicated possible binding and modification of FMN due to the formation of PhtDa:FMN:DMAP complex. UV-VIS measurement of purified PhtDb after incubation with FMN and DMAP showed absorption peak maxima at 384 nm and 458 nm (Figure 4B), which are the characteristic peaks for the FMN molecule (Lin *et al*., 2015). However, slight changes were observed in PhtD:FMN spectrum that consisted of a decrease in the absorbance intensity at 380 nm and ̴ 450 nm and the developed flat shoulders at 380 and 450 nm (Figure 4B). This strongly indicates an interaction of FMN with the PhtDb.

In addition to this, structural modelling of both PhtDa and PhtDb suggested that only PhtDb possesses the FMN-binding domain and forms a dodecameric complex (264 kDa; 12 FMN per 12 PhtDb), whereas PhtDa is a homodimer based on the known template structure (Rangarajan *et al*., 2004; (Figure 7B and 5C)). Superimposition of the model structure of both PhtDa and PhtDb with the respective templates of known crystal structures such as homodimer Pad1 in *P. aeruginosa* ((PDB code 4iws); Jacewicz *et al*., 2013) showed high overlapping and similarity in the overall fold of the secondary structure elements with the predicted model of PhtDa (Figure 7C). Whereas superimposition of the homo 12-mer crystal structure of FMN-dependent UbiX-like enzyme (PDB code 1sbz) in *E. coli* O157:H7 (Rangarajan *et al*., 2004) exhibited very high overlapping of predicted homo-12mer 3-D model of PhtDb (Figure 7D). Therefore, this accurately reflects that subunits in both PhtDa and PhtDb are arranged in a similar pattern with the oligomeric structures of their respective templates and thus it is most likely to have functional homology (Figure 7).

Native protein analysis of cell-free extract of *Azoarcus* sp strain PA01 as well as purified PhtDa and PhtDb revealed the formation of a protein complex of the expected molecular weight in analogy to the oligomeric structure of the related proteins as described in Figure 7. Based on the 3-D structure modelling, PhtDa (60 kDa) forms a 120 kDa homodimer while the PhtDb (22 kDa) is homo dodecamer forming a 264 kDa oligomer complex. Thus, both together (PhtDa and PhtDb) are expected to form a 384 kDa (approx. 390 kDa) protein complex. Native protein analysis of cell-free extract of *Azoarcus* sp. strain PA01 cells grown in *o*-phthalate showed the protein bands, particularly at approximate 380 and 350 kDa size (Figure 8A). MS analysis results (Table 1) of these two bands (2 and 3) surprisingly showed the identification of mainly PhtDa and PhtDb, with significant sequence coverage and score (Table 1) among the other proteins detected (data not shown). However, MS identification of the following lower molecular weight size protein bands (between 140 to 80 kDa), resulted in the identification of mainly PhtSa (43 kDa) and PhtSb (41 kDa), the expected subunits of the succinyl-CoA dependent CoA-transferase that is involved in the activation of *o*-phthalate to *o*-phthalyl-CoA (Junghare *et al*., 2016). The CoA-transferase are expected to form a tetramer (approximately 170 kDa) homologous to succinyl-CoA:*(R)*-benzylsuccinate CoA-transferase involved in toluene activation by *T. aromatica* (Leutwein and Heider, 2001). In addition to this, native protein analysis of partially purified PhDb incubated with FMN plus DMAP showed a single protein band at 390 kDa size, although the native protein analysis of PhtDa and PhtDb individually formed two distinct protein bands of 400 to 350 kDa (Figure 8B). Yet, the SDS-PAGE analysis of the purified PhtDa and PhtDb exhibited single bands at the expected sizes of 60 and 22 kDa (Figure 2C).

In conclusion, our experiments demonstrate that PhtDa and PhtDb constitute a two-component decarboxylase system for *o*-phthalyl-CoA decarboxylation similar to the UbiX and UbiD two-component decarboxylase system in which UbiX with its prenylated FMN cofactor is essential for the decarboxylase UbiD (Lin *et al*., 2015; White *et al*., 2015). Therefore, we strongly suggest that the prenylated FMN consequently plays a crucial role in the decarboxylation of *o*-phthalyl-CoA to benzoyl-CoA. However, how exactly prenylated FMN facilitates the decarboxylation of *o*-phthalyl-CoA to benzoyl-CoA is yet to be established.

## Materials and methods

### Materials and reagents

All chemicals used in this study were of analytical grade. Coenzyme A (CoA) trilithium salt, ampicillin, flavin mononucleotide (FMN), flavin dinucleotide (FAD), γ,γ-dimethylallyl monophosphate (DMAP), phthalate isomers and phthalic anhydride were from Sigma-Aldrich (Germany. *o*-Phthalyl-CoA’s used in this study were produced as described previously (Junghare *et al*., 2016). Materials and equipment for protein purification were obtained from GE Healthcare (Munich, Germany).

### Strains and cultivation conditions

*Azoarcus* sp. strain PA01 was cultivated anoxically in non-reduced, freshwater mineral medium containing *o*-phthalate (2 mM) or benzoate (2 mM) as the growth substrate supplemented with 10 - 12 mM nitrate as electron acceptor as described previously (Junghare *et al*., 2015b; Junghare *et al*., 2016). Cells were harvested by centrifugation (10,000 x g for 10 min at 10°C) at an optical density (OD_600nm_) 0.2 - 0.3 and stored at −20 °C until further use. *E. coli* OneShot TOP10, *E. coli* BL21 star (DE3) and *E. coli* Rosetta 2 (DE3)pLysS (Invitrogen) cells were grown either in Luria Bertani (LB) (Sambrook *et al*., 1989) or in terrific broth (TB) (Tartoff *et al*., 1987) media (unless otherwise mentioned) supplemented with 100 µg/ml of ampicillin.

### Gene amplifications, plasmid construction and sequencing

The genes encoding *o*-phthalyl-CoA decarboxylase, i.e., PA01_00217 (1584 bp) and PA01_00218 (600 bp) sequence can be found at the Integrated Microbial Genomes (IMG) under gene ID 2597201524 and 2597201525, (https://img.jgi.doe.gov/cgi-bin/m/main.cgi) respectively, in the draft genome sequence of *Azoarcus* sp. strain PA01 (Junghare *et al*., 2015b). For PCR amplification of the respective protein (PhtDa and PhtDb) coding genes, primers were synthesized (Microsynth, Switzerland). To facilitate directional cloning of the genes into champion pET100/D-TOPO expression vector which introduces a C-terminal 6x-His-tag (3 kDa), overhang of four nucleotides (CACC as underlined) was added to 5’ end of each forward primer of the respective genes. The gene PA01_00217 coding for PhtDa (60 kDa) was amplified using the primers: 217f, 5’-<u>CACC</u>ATGAACGATCTGGCAACGA-3’ and 217r, 5’-CTTTGTGGTGGCAGCAGC-3’, and PA01_00217 coding for PhtDb (22 kDa) was amplified using primers: 218f, 5’-<u>CACC</u>ATGAAAAGACTGATAGTGGGG-3’ and 218r, 5’-TGCGATTACTTCTGCGGC-3’, respectively. Gene amplification was performed using proof reading Phusion High Fidelity (HF) DNA polymerase (Finnzymes) in the PCR reaction (50 µl) containing: 2 µl of gDNA template (approx. 50 −100 ng); 2 µl of each primer (100 µM); 1 µl of 50 mM MgCl_2_ (unless otherwise mentioned); 10 µl of 5x Phusion HF buffer; 20 µl of dNTPs (500 µM); 0.5 µl of Phusion HF DNA polymerase (0.25 Units) and 12.5 µl of molecular grade water (for amplification of gene PA01_00218, MgCl_2_ was not added). A personal DNA thermal cycler (Eppendorf) was used for PCR reactions: initial denaturation at 98 °C for 5 min followed by 35 cycles of 98 °C for 30 s, 65 °C (70 °C for PA01_00217) for 40 s, 72 °C for 90 s and final extension at 72 °C for 10 min. PCR reactions were analysed by agarose (1 % w/v) gel electrophoresis and stained with ethidium bromide. Analysed PCR products were directly purified using DNA Clean and Concentrator kit (Zymo Research). DNA concentrations were determined using a nano UV spectrophotometer (Thermo Scientific).

The purified PCR products were cloned by mixing with the champion pET100/D-TOPO expression vector and ligated according to the manufacturer’s protocol (Invitrogen). The ligated vector containing the gene of interest (recombinant plasmid) was transformed into chemically competent *E. coli* OneShot TOP10 cells and grown on LB agar plates containing ampicillin (100 µg/ml). Positive clones containing the gene of interest were screened using the T7 primer pair as: T7, 5’-TAATACGACTCACTATAGGG-3’ and T7 reverse, 5’-TAGTTATTGCTCAGCGGTGG-3’, provided in the champion directional TOPO expression kit. Positive clones of *E. coli* TOP10 cells were propagated in LB medium (ampicillin, 100 µg/ml) and used for recombinant plasmid DNA isolation (Zypy Plasmid Miniprep, Zymo Research). Prior to plasmid transformation to expression host *E. coli* cells, the correct integration and sequence identity of the inserted gene was individually verified by sequencing the recombinant plasmids across the cloning site using the T7 primers (as mentioned above). DNA sequencing was performed at GATC-Biotech (Constance, Germany) and sequences were compared with the original gene sequences from the genome of *Azoarcus* sp. strain PA01 (Junghare *et al*., 2015b) for any mutation or base change using the BLAST (http://www.ncbi.nlm.nih.gov).

### Optimization of protein expression and purifications

Chemically competent OneShot *E. coli* BL21 Star (DE3) and *E. coli* Rosetta 2 (DE3)pLysS cells were transformed with the purified recombinant vector. Transformant’s were grown either in 10 ml of LB liquid or onto the LB solid agar medium (for maintenance) containing ampicillin (100 µg/ml). Over-expression of recombinant proteins PhtDa and PhtDb were performed in LB medium or in TB medium containing 100 µg/ml of ampicillin supplemented with 1 % glucose and 3 % ethanol (unless otherwise mentioned). Pre-cultures were grown initially at 37 °C under shaking conditions (200 rpm). Cells were induced for protein expression at an optical density (OD_600nm_) between 0.5 to 0.6 by adding 0.3 - 0.5 mM of isopropyl-β-D-thiogalactopyranoside (IPTG) and grown at different temperatures (at 37 °C for about 4 h; 25 °C for about 7 h and at 15 °C for about 17 h). Cells were harvested by centrifugation (10,000 x g for 10 min at 8 °C) and stored frozen at −20 °C until further use or analysed for soluble protein expression. Protein expression optimization experiments were performed in small scale cultures using 10 to 100 ml of growth media.

For large scale protein expression, transformed cultures were grown in 500 or 1000 ml media. About 2 to 3 g wet cell material was suspended in 5 ml lysis buffer containing 50 mM Tris-HCl (pH 7.6), 200 mM NaCl, 100 mM KCl, 60 mM imidazole, 10 % glycerol and 5 mM dithiotritol designated as buffer A. Prior to cell lysis, the cell suspension was supplemented with 1 mg/ml lysozyme (Sigma), 0.01 mg/ml DNase I (Sigma), 50 mM L-glutamate, 50 mM L-arginine-HCl, 1 mg of protease inhibitor cocktail (Roche) plus dimethyl allyl monophosphate (DMAP) together with 1 mM flavin mononucleotide (FMN) or 1 mM flavin dinucleotide (FAD). The cell suspension was incubated on ice for about 30 min for enzymatic (lysozyme) cell lysis. Finally, cells were passed through a French pressure cell operated at 139 kPa at least for three times. Cell lysate was centrifuged at 27,000 x g for 30 min at 4 °C to remove the cell debris. The supernatant (cell-free extract) was either used directly for enzyme activity measurements or subjected to further enzyme purification.

His-tagged recombinant enzyme purification was performed using the Ni^2+^ ion affinity chromatography. *E. coli* supernatant was directly applied onto a 1-ml Ni^2+^ ion chelating poly-histidine affinity column (HisTrap HP, GE Healthcare) equilibrated with buffer A (50 mM Tris-HCl, pH 7.6; 200 mM NaCl; 100 mM KCl; 60 mM imidazole; 10 % glycerol and 5 mM dithiotritol). After washing the column with 10 ml of buffer A, the His-tagged proteins were finally eluted with 2 - 3 ml of elution buffer B (buffer A supplemented with 0.5 M imidazole final concentration). Imidazole was removed by passing the eluted protein fraction over a desalting NAP^TM^-25 column (GE Healthcare) equilibrated with buffer A and eluted with 2.5 ml of the same buffer. Fifty percent glycerol was added to the purified protein to a final concentration of 25 % (v/v) and stored at −20 °C. Protein contents were determined using the Bradford assay (using BAS as the standard).

### Molecular weight determination by SDS-PAGE

Sodium dodecyl sulfate-polyacrylamide gel electrophoresis (SDS-PAGE) analysis was used to determine the purity and molecular weight of the over-expressed proteins. SDS-PAGE was also used to determine the expression level of soluble protein in *E. coli* cell-free extracts or cell pellet, according to Laemeli method using the established protocol (Junghare *et al*., 2016). Protein samples were mixed with sample loading buffer (0.05 % bromophenol blue, 5 % β-mercaptoethanol, 10 % glycerol, 1 % SDS in 0.25M Tris-HCl buffer pH 6.8) and denatured by heating at 99 °C for 10 min. Samples were loaded onto the one-dimensional SDS gel (12 % resolving and 4 % stacking gel) and electrophoresis was performed using a BioRad Mini-Protean cell (8 cm x 6 cm x 1.0 mm) at constant voltage of 120 V for about 2 h. For molecular weight (MW) estimation of protein bands, 10 µl of the prestained protein ladder of MW range 10 - 180 kDa was used (Page Ruler, Thermo Fisher). Protein bands were visualised with 0.25 % Coomassie Brilliant Blue R-250 dye dissolved in 10 % acetic acid and 50 % methanol, followed by de-staining in 10 % acetic acid and 50 % methanol in distilled water. The detected protein band size was compared with standard protein MW markers for size estimation (Pre-stained protein molecular size marker 15 – 100 kDa, Themo Scientific). Expected size protein bands were excised from the SDS gel and given for peptide identification by electrospray mass spectrometry (MS) analysis at the Proteomics Facility, University of Konstanz. The MS spectra of peptides were analysed using the Mascot search engine (version 2.2.03, Matrix Science) to confirm the identity of the over-expressed proteins by comparison with the deduced amino acid sequence of the respective genes from the genome of the *Azoarcus* sp. strain PA01 (Junghare *et al*., 2015b).

### Decarboxylase activity assays and cofactor requirement

*o*-Phthalyl-CoA decarboxylase activity of PhtDa and PhtDb enzymes overexpressed in *E. coli* cells was assayed according method described before (Junghare *et al*., 2016). The decarboxylase activity was assayed directly with the *E. coli* cell lysate containing overexpressed protein PhtDa and PhtDb for conversion of *o*-phthalyl-CoA to benzoyl-CoA. Standard anoxic enzyme assays (150 µl) contained: 100 µl tri-ethanolamine buffer (0.1M, pH 7.6), 10 - 20 µl cell lysates (PhtDa and PhtDb) and 1 mM dithionite. Enzyme reaction was started by the addition of 10 µl *o*-phthalyl-CoA and incubated at 30 °C for about 30 min. Samples (30 µl) were removed at different time intervals and proteins were precipitated with methanol (60 µl). Samples were centrifuged and the supernatants were analysed for benzoyl-CoA formation by liquid chromatography coupled with electrospray ionization-MS (LC-ESI-MS) as described previously (Junghare *et al*., 2016). The substrate specificity of the decarboxylase activity was assayed for decarboxylation of isophthalyl-CoA, terephthalyl-CoA or fluoro-*o*-phthalyl-CoA to benzoyl-CoA (fluoro-benzoyl-CoA in case fluoro-*o*-phthalyl-CoA). Effect of oxygen exposure on the stability and activity of PhtDa and PhtDb was assayed by performing enzyme assay under oxic condition.

The flavin dependent activity of recombinant PhtDb (PA01_00218) was determined using reconstitution experiments. Heterologously expressed PhtDb in *E. coli* cells were suspended in cell lysis buffer A (as mentioned above) and supplemented with 1 mM flavin dinucleotide (FAD) or flavin mononucleotide (FMN) plus about 10 µl of di-methyl allyl mono phosphate (DMAP). Cell were incubated on ice for about 30 min (enzymatic cell lysis) and finally opened by French Press treatment as described above. Cell lysate was either used directly for enzyme activity determinations or processed further by centrifugation (27, 000 x g for 30 min at 4 °C) for His-tag purification of recombinant protein. UV-visible spectrum (300nm to 500nm wavelength) of the purified PhtDb enzyme was recorded in the UV-vis spectrophotometer (Jasco V-630, spectrophotometer) for determination of FMN content. Spectrum of 1 µM FMN in elution buffer was used as reference.

### Native protein PAGE analysis

Estimation of protein complex size and protein-protein interaction, native protein polyacrylamide gel electrophoresis (PAGE) was performed with the cell-free extract of *Azoarcus* sp. strain PA01 cells grown with *o*-phthalate and benzoate as the control. Additionally, native protein gel was also performed with the partially His-tag purified recombinant PhtDa (60 kDa) and PhtDb (22 kDa) proteins. Approximately, each of 30 µg of purified PhtDa and PhtDb proteins or soluble proteins from the cell-free extract of *Azoarcus* sp. strain PA01 cells were mixed with the native protein sample loading buffer (60 mM Tris-HCl, pH 6.8; 30 % glycerol; 0.01 % (w/v) bromophenol blue). PhtDa and PhtDb protein-protein interaction were investigated by incubating these two proteins together plus FMN and DMAP addition. Electrophoresis was performed using BioRad Protean Mini cell and the PreCast gel slab (BioRad) at 130 V for about 3 h. Protein bands were visualised by straining the gels with Coomassie Brilliant Blue R-250 as described above. Protein-protein interaction and protein complex sizes were estimated by comparing the stained protein bands with the native protein molecular weight markers range between 60 to 690 kDa (Amersham, GE Healthcare Life Sciences). Protein bands excised from the stained native protein gels were identified by MS analysis.

### Sequence alignment and structure homology modelling

Protein similarity searches were performed using the BLASTp search tool available at the NCBI web page (http://www.ncbi.nlm.nih.gov/BLAST/). A phylogenetic tree was constructed to determine sequence divergence and the evolutionary positions using the MEGA 7 software package (Kumar *et al*., 2016). Amino acid sequences of the PhtDb and PhtDa of *Azoarcus* sp. strain PA01 and the characterized the UbiX-like and UbiD-like enzymes from bacterial and fungal species were considered for tree construction. Phylogenetic tree was calculated using neighbor-joining method and the bootstrap (1000 replicates) method was followed for corroborating the consistency of each node in the tree. For determination of conserved amino acid regions, the T-Coffee Expresso multiple sequence alignment package with the CLUSTAL W (1.83) alignment tool (http://tcoffee.crg.cat/apps/tcoffee/do:expresso) was used (Notredame *et al*., 2000). Aligned sequences were edited in the ESPript 3.0 program ((http://espript.ibcp.fr/ESPript/ESPript/index.php); Robert and Gouet, 2014). Homology-based 3-dimensional (3-D) models were constructed for PhtDa and PhtDb proteins to understand structural aspects and functional homology to the related UbiD-like and UbiX-like enzymes. The predicted structural models for PhtDa and PhtDb were generated through the SWISS-MODEL workspace in an automated mode (Arnold *et al*., 2006; Biasini *et al*., 2014). PhtDa structure was predicted using the template putative aromatic acid decarboxylase (Pad1) crystal structure (PDB accession code 4iws.1.A) from *Pseudomonas aeruginosa* (Jacewicz *et al*., 2013), whereas PhtDb structure was predicted using the FMN-dependent Ubix-like decarboxylase (PDB code 1sbz.1.A) from *Escherichia coli* O157:H7 (Rangarajan *et al*., 2004). Predicted structural model visualization and superimpositions analyses with the respective template structures were performed with the UCSF Chimera package (Pettersen *et al*., 2004).

## Acknowledgments

MJ acknowledges the University of Konstanz for financial support. MJ greatly acknowledges Jasmin Frey for helpful suggestions and discussions during the cloning and heterologous protein expression experiments. MJ greatly thanks to Prof. Dieter Spiteller and Prof. Bernhard Schink for reading and improvement of the manuscript.

## Notes

### Competing Interest Statement

The authors have declared no competing interest.

## References

Aftring, R. P., Chalker, B. E., Taylor, B. F. (1981). Degradation of phthalic acids by denitrifying, mixed cultures of bacteria. Appl Environ Microbiol 41: 1177–1183.

Arnold, K., Bordoli, L., Kopp, J., Schwede, T. (2006). The SWISS-MODEL Workspace: A web-based environment for protein structure homology modelling. Bioinformatics, 22,195–201.

Baba, T., Ara, T., Hasegawa, M., Takai, Y., Okumura, Y., et al. (2006). Construction of *Escherichia coli* K-12 in-frame, single-gene knockout mutants: the Keio collection. Mol Syst Biol 2: 2006–0008.

Batie, C. J., LaHaie, E., Ballou, D. (1987). Purification and characterization of phthalate oxygenase and phthalate oxygenase reductase from *Pseudomonas cepacia*. J Biolog Chem. 262: 1510–1518.

Battersby, N. S., Wilson, V. (1989). Survey of the anaerobic biodegradation potential of organic chemicals in digesting sludge. Appl Environ Microbiol 55: 433–439.

Beller, H. R., Spormann, A. M., Sharma, P. K. (1996). Isolation and characterization of a novel toluene degrading sulfate reducing bacterium. Appl Environ Microbiol 62: 1188–1196.

Bemis, A. G., Dindorf, J. A., Horwood, B., Samans, C. (1982). Phthalic acids and other benzenepolycarboxylic acids. in Kirk Othmer encyclopedia of chemical technology, eds Mark H. F., Othmer D. F., Overberg C. G., Seaborg G. T., Grayson M., Eckroth D. (John Wiley & Sons, New York, N.Y), 17: 732–777.

Bhuiya, M. W., Lee, S. G., Jez, J. M. (2015). Structure and mechanism of ferulic acid decarboxylase (FDC1) from *Saccharomyces cerevisiae*. Appl Environ Microbiol 81: 1216–1223.

Biasini, M., Bienert, S., Waterhouse, A., Arnold, K., Studer, G., Schmidt, T., Kiefer, F., Cassarino, T. G., Bertoni, M., Bordoli, L., Schwede, T., (2014). SWISS-MODEL: modelling protein tertiary and quaternary structure using evolutionary information. Nucleic Acids Research 42: W252–W258.

Boll, M. (2005a). Key enzymes in the anaerobic aromatic metabolism catalysing Birch-like reductions. Biochim Biophys Acta 1707: 34–50.

Boll, M. (2005b). Dearomatizing benzene ring reductases. J Mol Microbiol Biotechnol 10: 132–142.

Boll, M., Fuchs, G. (1995). Benzoyl-coenzyme A reductase (dearomatizing), a key enzyme of anaerobic aromatic metabolism. ATP dependence of the reaction, purification and some properties of the enzyme from *Thauera aromatica* strain K172. Eur J Biochem 234: 921–933.

Boll, M., Fuchs, G., Heider, J. (2002). Anaerobic oxidation of aromatic compounds and hydrocarbons. Curr Opin Chem Biol 6: 604–611.

Boll, M., Laempe, D., Eisenreich, W., Bacher, A., Mittelberger, T., Heinze. J., Fuchs, G. (2000). Non-aromatic products from anoxic conversion of benzoyl-CoA with benzoyl-CoA reductase and cyclohexa-1,5-diene-1-carbonyl-CoA hydratase. Biol Chem 275: 21889–21895.

Boonyaroj, V., Chiemchaisri, C., Chiemchaisri, W., Theepharaksapan, S., Yamamoto, K., (2012). Toxic organic micro-pollutants removal mechanisms in long-term operated membrane bioreactor treating municipal solid waste leachate. Bioresour Technol 113: 174–180.

Breese, K., Boll, M., Alt-Morbe, J., Schagger, H., Fuchs, G. (1998). Genes coding for the benzoyl-CoA pathway of anaerobic aromatic metabolism in the bacterium *Thauera aromatica*. Eur J Biochem 256: 148–154.

Buckel, W., Keese, R. (1995). One electron reactions of CoASH esters in anaerobic bacteria. Angew Chem Int Ed Engl 34: 1502–1506.

Camacho-Munoz, D., Martin, J., Santos, J. L., Alonso, E., Aparicio, I., De la Torre, T., et al., (2012). Effectiveness of three configurations of membrane bioreactors on the removal of priority and emergent organic compounds from wastewater: comparison with conventional wastewater treatments. J Environ Monit 14: 1428–1436.

Carmona, M., Zamarro, M. T., Blázquez, B., Durante-Rodríguez, G., Juárez, J. F., Valderrama, J. A., Barragán, M. J., García, J. L., Díaz, E. (2009). Anaerobic catabolism of aromatic compounds: a genetic and genomic view. Microbiol Mol Biol Rev 73: 71–133.

Carrara, S., Morita, D. M., Boscov, M. E. G. (2011). Biodegradation of di(2-ethylhexyl)phthalate in a typical tropical soil. J Hazard Mater 197: 40–48.

Cashion, P., Hodler-Franklin, M.A., McCully, J., Franklin, M. (1977). A rapid method for base ratio determination of bacterial DNA. Anal Biochem 81: 461–466.

Castro, H. F., Williams, N. H., Ogram, A. (2000). Phylogeny of sulfate reducing bacteria. FEMS Microbiol Ecol 31: 1–9.

Chang, B. V., Liao, C. S., Yuan, S. Y. (2005). Anaerobic degradation of diethyl phthalate, di-nbutyl phthalate, and di-(2-ethylhexyl) phthalate from river sediment in Taiwan. Chemosphere 58: 1601–1607.

Chang, H. K., Zylstra, G. J. (1998). Novel organization of the genes for phthalate degradation from *Burkholderia cepacia* DBO1. J Bacteriol 180: 6529–6537.

Chao, W. L., Lin, C. M., Shiung, I. I., Kuo, Y. L. (2006). Degradation of of di-butyl-phthalate by soil bacteria. Chemosphere 63: 1377–1383.

Chen, J. A., Li, X., Li, J., Cao, J., Qiu, Z. Q., Zhao, Q., Xu, C., Shu, W. Q. (2007). Degradation of environmental endocrine disruptor di-2-ethylhexyl phthalate by a newly discovered bacterium, *Microbacterium* sp. strain CQ0110Y. Appl Microbiol Biotechnol 74: 676–682.

Chen, M. H., Sheu, S. Y., James, E. K., Young, C. C., Chen, W. M. (2013). *Azoarcus olearius* sp. nov., a nitrogen-fixing bacterium isolated from oil-contaminated soil. Int J Syst Evo Microbiol 63: 3755–3761.

Cheung, J. K. H., Lam, R. K. W., Shi, M. Y., Gu, J. D. (2007). Environmental fate of endocrine-disrupting dimethyl phthalate esters (DMPE) under sulfate-reducing condition. Sci Total Envion 381: 126–133.

Dagley, S. (1978). Determinants of biodegradability. Q Rev Biophys 11: 577–602.

Dagley, S., Chapman, P. J., Gibson, D. T., Wood, J. M. (1964). Degradation of benzene nucleus by bacteria. Nature 202: 775–778.

Dagley, S., Evans, W. C., Ribbons, D. W. (1960). New pathways in the oxidative metabolism of aromatic compounds by microorganisms. Nature 188: 560–566.

Eaton, R. W. (2001). Plasmid-encoded phthalate catabolic pathway in *Arthrobacter keyseri* 12B. J Bacteriol 183: 3689–3703.

Eaton, R. W., Ribbons, D. W. (1982). Metabolism of dimethylphthalate by *Micrococcus* sp. strain 12B. J Bacteriol 151: 465–467.

Ebenau-Jehle, C., Mergelsberg, M., Fischer, S., Brüls, T., Jehmlich, N., von Bergen, M., Boll, M. (2016). An unusual strategy for the anoxic biodegradation of phthalate. The ISME Journal doi:10.1038/ismej.2016.91.

Ejlertsson, J., Svensson, B. H. (1996). Degradation of bis(2-ethylhexyl) phthalate constituents under methanogenic conditions. Biodegradation 7: 501–506.

Engelhardt, G., Wallnofer, P. R., Hutzinger. (1975). The microbial metabolism of di-n-butyl phthalate and related dialkyl phthalates. Bull Environ Contamin Toxicol 13: 342–347.

Evans, W. C. (1977). Biochemistry of the bacterial catabolism of aromatic compounds in anaerobic environments. Nature 270: 17–22.

Evans, W. C., Fuchs, G. (1988). Anaerobic degradation of aromatic compounds. Annu Rev Microbiol 42: 289–317.

Fang, H. H. P., Chen, T., Li, Y. Y., Chui, H. K. (1996). Degradation of phenol in wastewater in an upflow anaerobic sludge blanket reactor. Water Resources 30: 1353–1360.

Fatoki, O. S., Vernon, F. (1990). Phthalate esters in rivers of the Greater Manchester area, U.K. Sci Total Environ 95: 227–232.

Felsenstein, J. (1981). Evolutionary trees from DNA sequences: a maximum likelihood approach. J Mol Evol 17: 368–376.

Field D, Garrity G, Gray T, Morrison N, Selengut J, Sterk P, Tatusova T, Thomson N, Allen MJ, Angiuoli SV, et al. (2008). The minimum information about a genome sequence (MIGS) specification. Nat Biotechnol 26: 541–547.

Fitch, W. M. (1971). Toward defining the course of 260 evolution: minimum change for a specific tree topology. Syst Zool 20: 406–416.

Foght, J. (2008). Anaerobic biodegradation of aromatic hydrocarbons: pathways and prospects. J Mol Microbiol Biotechnol 15: 93–120.

Forward, J. A., Behrendt, M. C., Wyborn, N. R., Cross, R., Kelly, D. J. (1997). TRAP transporters: a new family of periplasmic solute transport systems encoded by the dctPQM genes of *Rhodobacter capsulatus* and by homologs in diverse gram-negative bacteria. J Bacteriol 179: 5482–5493.

Friedrich, M., Springer, N., Ludwig, W., Schink, B. (1996). Phylogenetic positions of *Desulfofustis glycolicus* gen. nov., sp. nov., and *Syntrophobotulus glycolicus* gen. nov., sp. nov., two new strict anaerobes growing with glycolic acid. Int J Syst Evol Microbiol 46: 1065–1069.

Fritsche, W., Hofrichter, M. (2008). Aerobic degradation by microorganisms. In: Environmental processes-soil decontamination, J Klein, Ed., Wiley-VCH, Weinheim, Germany. p. 146–155.

Fuchs, G. (2008). Anaerobic metabolism of aromatic compounds. Ann N Y Acad Sci 1125: 82–99.

Fuchs, G., Boll, M., Heider, J. (2012). Microbial degradation of aromatic compounds - from one strategy to four. Nat Review Microbiol 9: 803–816.

Fuchs, G., Mohamed, M. E. S., Altenschmidt, U., Koch, J., Lack, A., Brackmann, R., Lochmeyer, C., Oswald, B. (1994). Biochemistry of anaerobic biodegradation of aromatic compounds. In Biochemistry of microbial degradation. ed Ratledge C. (Kluwer Academic Publishers, Dordrecht, The Netherlands), p. 513-553.

Fukuhara, Y., Inakazu, K., Kodama, N., Kamimura, N., Kasai, D., Katayama, Y., Fukuda, M., Masai, E. (2010). Characterization of the isophthalate degradation genes of *Comamonas* sp. strain E6. Appl Environ Microbiol 76: 519–527.

Fukuhara, Y., Kasai, D., Katayama, Y., Fukuda, M., Masai, E. (2008). Enzymatic properties of terephthalate 1,2-dioxygenase of *Comamonas* sp. strain E6. Biosci. Biotechnol. Biochem. 72: 2335–2341.

Fushiwaki, Y., Niino, T., Ishibashi, T., Takeda, K., Onodera, S. (2003). Tumor-promoting activity of phthalate esters estimated by in vitro transformation using Bhas cells. J Health Sci 49: 82–87.

Gao, D. W., Wen, Z. D. (2016). Phthalate esters in the environment: A critical review of their occurrence, biodegradation, and removal during wastewater treatment processes. Sci Total Environ 541: 986–1001.

Garrity, G. M., Bell, J. A., Lilburn, T. Class II. Betaproteobacteria class. nov. In: Garrity GM, Brenner DJ, Krieg NR, Staley JT (eds), Bergey’s Manual of Systematic Bacteriology, Second Edi-tion, Volume 2, Part C, Springer, New York, 2005b, p. 575.

Garrity, G. M., Bell, J. A., Lilburn, T. Family I. *Rhodocyclaceae* fam. nov., In: DJ Brenner, NR Krieg, JT Staley, (eds) GG (eds), Bergey’s Manual of Systematic Bacteriology, Second Edition, Vol-ume 2, Part C, Springer, New York, 2005d, p. 887.

Garrity, G. M., Bell, J. A., Lilburn, T. Order VI. *Rhodocyclales* ord. nov. In: Garrity GM, Brenner DJ, Krieg NR, Staley JT (eds), Bergey’s Manual of Systematic Bacteriology, Second Edition, Volume 2, Part C, Springer, New York, 2005c, p. 887.

Garrity, G. M., Bell, J. A., Lilburn, T., Phylum XIV. *Proteobacteria* phyl. In: Garrity GM, Brenner DJ, Krieg NR, Staley JT, editors. Bergey’s Manual of Systematic Bacteriology, Second Edition, Volume 2, Part B. 2nd ed. New York: Springer; 2005a.

Giam, C. S., Chan, H. S., Neff, G. S., Atlas, E. (1978). Phthalate ester plasticizers: A new class of marine pollutant. Science 199: 419–421.

Giam, C.S., Atlas, E., Powers, M.A Jr., and Leonard, J.E. (1984). Phthalic acid esters. In: Hutzinger, O., ed., The handbook of environmental chemistry, vol. 3, part c, anthropogenic compounds, Berlin, Springer Verlag, p. 67–142.

Gibson, D. T., Parales, R, E. (2000). Aromatic hydrocarbon dioxygenases in environmental biotechnology. Curr Opin Biotechnol 11: 236–243.

Gibson, D. T., Subramanian, V. (1984). Microbial degradation of aromatic hydrocarbons. In: Gibson DT (ed) Microbial degradation of organic compounds. Marcel Dekker, Inc., New York, pp 181–252.

Gibson, G., Harwood, CS. (2002). Metabolic diversity in aromatic compound utilization by anaerobic microbes. Annu Rev Microbiol 56: 345–369.

Gittel, A., Seidel, M., Kuever, J., Galushko, A. S., Cypionka, H., Könneke, M. (2010). *Desulfopila inferna* sp. nov., a sulfate-reducing bacterium isolated from the subsurface of a tidal sand-flat. Int J Syst Evol Microbiol 60: 1626–1630.

Gledhill, W. E., Kaley, R. G., Adams, W. J., Hicks, O., Michael, P. R., Saeger, V. W., et al., (1980). An environmental safety assessment of butyl benzyl phthalate. Environ Sci Technol 14: 301–305.

Gregersen, T. (1978). Rapid method for distinction of Gram-negative from Gram-positive bacteria. Eur J Appl Microbiol Biotechnol 5: 123–127.

Grifoll, M., Selifonov, S.A., Chapman P. J. (1994). Evidence for a novel pathway in the degradation of fluorene by *Pseudomonas* sp. strain F274. Appl Environ Microbiol 60: 2438–2449.

Gulmezian, M., Hyman, K. R., Marbois, B. N., Clarke, C. F., Javor, G. T. (2007). The role of UbiX in *Escherichia coli* coenzyme Q biosynthesis. Arch. Biochem. Biophys. 467, 144–153.

Habe, H., Miyakoshi, M., Chung, J., Kasuga, K., Yoshida, T., Nojiri, H., Omori, T. (2003). Phthalate catabolic gene cluster is linked to the angular dioxygenase gene in *Terrabacter* sp. strain DBF63. Appl Microbiol Biotechnol 61: 44–54.

Han, R. (2008). Phthalate biodegradation: gene organization, regulation and detection. PhD thesis, New Brunswick, New Jersey.

Harwood, C. S., Burchhardt, G., Herrmann, H., Fuchs, G. (1999). Anaerobic metabolism of aromatic compounds via the benzoyl-CoA pathway. FEMS Microbiol Rev 22: 439–458.

Harwood, C. S., Parales, R. E. (1996). The b-ketoadipate pathway and the biology of self-identity. Ann Rev Microbiol 50: 553–590.

Heider, J. (2001). A new family of CoA-transferases. FEBS Lett 509: 345–349.

Heider, J., Fuchs, G. (1997). Anaerobic metabolism of aromatic compounds. Eur J Biochem 243: 577–596.

Hyatt, D., Chen, G. L., Locascio, P. F., Land, M. L., Lar-imer, F. W., Hauser, L. J. (2010). Prodigal: prokaryotic gene recognition and translation initiation site identification. BMC Bioinformatics 11: 119.

Jacewicz, A., Izumi, A., Brunner, K., Schnell, R., Schneider, G. (2013). Structural insights into the UbiD protein family from the crystal structure of PA0254 from *Pseudomonas aeruginosa*. PLoS ONE 8: 63161.

Janssen, P. H., Schuhmann, A., Bak, F., Liesack, W. (1996). Disproponation of inorganic sulfur compounds by the sulfate reducing bacterium *Desulfocapsa thiozymogenes* gen. nov., sp. nov. Arch Microbiol 166: 184–192.

Jonsson, S., Vavilin, V., Svensson, B. (2006). Phthalate hydrolysis under landfill conditions. Water Sci Technol 53: 119–127.

Junghare, M., Patil, Y., Schink, B. (2015b). Draft genome sequence of a nitrate-reducing, o-phthalate degrading bacterium, *Azoarcus* sp. strain PA01. Stand Genomic Sci 10: 90.

Junghare, M., Schink, B. (2015a). *Desulfoprunum benzoelyticum* gen. nov., sp. nov., a Gram-stain-negative, benzoate-degrading, sulfate-reducing bacterium isolated from a wastewater treatment plant. International Journal of Systematics and Evolutionary Microbiology 65: 77–84.

Junghare, M., Spiteller, D., Schink, B. (2016). Enzymes involved in the anaerobic degradation of *ortho*-phthalate by the nitrate-reducing bacterium *Azoarcus* sp. strain PA01. Environmental Microbiology 18: 3175–3188.

Junghare, M., Subudhi, S., Lal, B. (2012). Improvement of hydrogen production under decreased partial pressure by newly isolated alkaline tolerant anaerobe, *Clostridium butyricum* TM-9A: Optimization of process parameters. Int J of Hydrogen Energy 37: 3160–3168.

Kämpfer, P., Kroppenstedt, R. M. (1996). Numerical analysis of fatty acid patterns of coryneform bacteria and related taxa. Can J Microbiol 42: 989–1005.

Keyser, P., Pujar, B. G., Eaton, R. W., Ribbons, D. W. (1976). Biodegradation of the phthalates and their esters by bacteria. Environ Health Perspect 18: 159–166.

Kim, O. S., Cho, Y. J., Lee, K., Yoon, S. H., Kim, M., Na, H., Park, S. C., Jeon, Y. S., Lee, J. H., Yi, H., Won, S., Chun, J. (2012). Introducing EzTaxon: a prokaryotic 16S rRNA Gene sequence database with phylotypes that represent uncultured species. Int J Syst Evol Microbiol 62: 716–721.

Kiyohara, H., Nagao, K. (1978). The catabolism of phenanthrene and naphthalene by bacteria. J Gen Microbiol 105: 69–75.

Kleerebezem R, Pol LWH, Lettinga G (1999b) Anaerobic biodegradability of phthalic acid isomers and related compounds. Biodegradation 10:63–73.

Kleerebezem R, Pol LWH, Lettinga G (1999c) Energetics of product formation during anaerobic degradation of phthalate isomers and benzoate. FEMS Microbiol Ecology 29:273–282.

Kleerebezem, R., Hulshoff, P.L.W., Lettinga, G. (1999a). Anaerobic degradation of phthalate isomers by methanogenic consortia. Appl Environ Microbiol 65: 1152–1160.

Kopec, J., Schnell, R., Schneider, G. (2011). Structure of PA4019, a putative aromatic acid decarboxylase from *Pseudomonas aeruginosa*. Acta Crystallogr. F 67: 1184–1188.

Krause, A., Ramakumar, A., Bartels, D., Battistoni, F., Bekel, T., Boch, J., Böhm, M., Friedrich, F., Hurek, T., Krause, L., Linke, B., McHardy, A. C., Sarkar, A., Schneiker, S., Syed, A. A., Thauer, R., Vorhölter, F. J., Weidner, S., Pühler, A., Reinhold-Hurek, B., Kaiser, O., Goesmann, A. (2006). Complete genome of the mutualistic, N2-fixing grass endophyte *Azoarcus* sp. strain BH72. Nat Biotechnol 24: 1385–1391.

Kuever, J., Rainey, F. A., Widdel, F. (2005). Family II. *Desulfobulbaceae* fam. nov. In: D.J. Brenner, N.R. Krieg, J.T. Staley and G. M. Garrity. (editors), Bergey’s Manual of Systematic Bacteriology, second edition, vol. 2 (The Proteobacteria), part C (The Alpha-, Beta-, Delta-, and Epsilon proteobacteria), pp. 988–999. New York: Springer.

Kumar, S., Stecher, G., Tamura, K. (2016). MEGA7: molecular evolutionary genetics analysis version 7.0 for bigger datasets. Mol Biol Evol 33: 1870–1874.

Kung, J. W., Löffler, C., Dörner, K., Heintz, D., Gallien, S., Van Dorsselaer, A., et al. (2009). Identification and characterization of the tungsten-containing class of benzoyl-coenzyme A reductases. Proc Natl Acad Sci USA 106: 17687–17692.

Lambrot, R., Muczynski, V., Lécureuil, C., Angenard, G., Coffigny, H., Pairault, C. (2009). Phthalates impair germ cell development in the human fetal testis in vitro without change in testosterone production. Environ Health Perspect 117: 32–37.

Lane, D. J. (1991). 16S/23S rRNA sequencing: In Stackebrandt, E and Goodfellow, M (eds) Nucleic acid techniques in bacterial systematics. pp. 115-175. Chichester, England: John Wiley & Sons.

Latini, G., Del Vecchio A., Massaro, M., Verrotti, A., De Felice, C. (2006). Phthalate exposure and male infertility. Toxicology 226: 90–98.

Lau, T., Chu, W., Graham, N. (2005). The degradation of endocrine disruptor di-n-butyl phthalate by UV irradiation: a photolysis and product study. Chemosphere 60: 1045–1053.

Lee, D. J., Wong, B. T., Adav, S. S. (2014). *Azoarcus taiwanensis* sp. nov., a denitrifying species isolated from a hot spring. Appl Microbiol Biotechnol 98: 1301–1407.

Leutwein, C., Heider, J. (1999). Anaerobic toluene catabolic pathway in denitrifying *Thauera aromatica*: activation and b-oxidation of the first intermediate, (R)-(1)-benzylsuccinate. Microbiology 145: 3265–3271.

Leutwein, C., Heider, J. (2001). Succinyl-CoA:(R)-benzylsuccinate CoA-transferase: an enzyme of the anaerobic toluene catabolic pathway in denitrifying bacteria. J Bacteriol 183: 4288–4295.

Li, H., Gu, J. D. (2007). Complete degradation of dimethyl isophthalate requires the biochemical cooperation between *Klebsiella oxytoca* Sc and *Methylobacterium mesophilicum* Sr isolated from wetland sediment. Sci Total Environ 380: 181–187.

Li, J. X., Gu, J. D., Pan, L. (2005a). Transformation of dimethyl phthalate, dimethyl isophthalate and dimethyl terephthalate by *Rhodococcus rubber* Sa and modeling the processes using the modified Gompertz model. Int Biodeterior Biodegradation 55: 223–232.

Li, J. X., Gu, J. D., Yao, J. H. (2005b). Degradation of dimethyl terephthalate by *Pasteurella multocida* Sa and *Sphingomonas paucimobilis* Sy isolated from mangrove sediment. Int Biodeterior Biodegradation 56: 158–165.

Li, J., Chen, J. A., Zhao, Q., Li, X., Shu, W. Q. (2006). Bioremediation of environmental endocrine disruptor di-n-butyl phthalate ester by *Rhodococcus ruber*. Chemosphere 65: 1627–1633.

Li, Y. Y., Fang, H. H. P., Chen, T., Chui, H. (1995). UASB treatment of wastewater containing concentrated benzoate. J Environ Engineering 121: 748–751.

Liang, D., Zhang, T., Fang, H. H. P., He, J. (2008). Phthalates biodegradation in the environment. Appl Microbiol Biotechnol 80: 183–198.

Lie, T. J., Clawson, M. L., Godchaux, W., Leadbetter, E. R. (1999). Sulfidogenesis from 2-aminoethanesulfonate (taurine) fermentation by a morphologically unusual sulfate-reducing bacterium, Desulforhopalus singaporensis sp. nov. Appl Environ Microbiol 65: 3328–3334.

Lin, F., Ferguson, K. L., Boyer, D. R., Lin, X. N., Marsh, E. N. G. (2015). Isofunctional enzymes PAD1 and UbiX catalyze formation of a novel cofactor required by ferulic acid decarboxylase and 4-hydroxy-3-polyprenylbenzoic acid decarboxylase. ACS Chem Biol 10: 1137–1144.

Liolios, K., Mavromatis, K., Tavernarakis, N., Kyrpides, N. C. (2008). The Genomes On Line Database (GOLD) in 2007: status of genomic and metagenomic projects and their associated metadata. Nucleic Acids Res 36: 475–479.

Lipscomb, J. D. (2008). Mechanism of extradiol aromatic ring cleaving dioxygenases. Curr Opin Struct Biol 18: 644–649.

Liu, J., Liu, J. H. (2006). Ubiquinone (Coenzyme Q) biosynthesis in *Chlamydophila pneumoniae* AR39. Identification of the ubiD gene. Acta Biochim Biophys Sin 38: 725–730.

Lovley, D. R. (2003). Cleaning up with genomics: applying molecular biology to bioremidation. Nat Rev Microbiol 1: 35–44.

Lowe, T. M., Eddy, S. R. (1997). tRNAscan-SE: a program for improved detection of transfer RNA genes in genomic sequence. Nucleic Acids Res 25: 955–964.

Lykidis, A., Chen, C. L., Tringe, S. G., McHardy, A. C., Copeland, A., Kyrpides, N. C., Hugenholtz, P., Macarie, H., Olmos,A., Monroy, O., Liu, W. T. (2011). Multiple syntrophic interactions in a terephthalate-degrading methanogenic consortium. The ISME Journal 5: 122–130.

Markowitz, V. M., Mavromatis, K., Ivanova, N. N., Chen, I. M. A., Chu, K., Kyrpides, N. C. (2009). IMG-ER: a system for microbial genome annotation expert review and curation. Bioinformatics 25: 2271–2278.

Maruyama, K., Akita, K., Naitou, C., Yoshida, M., Kitamura, T. (2005). Purification and characterization of an esterase hydrolysing monoalkyl phthalates from *Micrococcus* sp. YGJ1. J Biochem 137: 27–32.

Matsumoto, M., Hirata-Koizumi, M., Ema, M. (2008). Potential adverse effects of phthalic acid esters on human health: a review of recent studies on reproduction. Regul Toxicol Pharm 50: 37–49.

Mavromatis, K., Ivanova, N. N., Chen, I. M., Szeto, E., Markowitz, V. M., Kyrpides, N. C. (2009). The DOE-JGI Standard operating procedure for the annotations of microbial genomes. Stand Genomic Sci 1: 63–67.

Mavromatis, K., Land, M. L., Brettin, T. S., Quest, D. J., Copeland, A., Clum, A., Goodwin, L., Woyke, T., Lapidus, A., Klenk, H. P., et al. (2012). The fast changing landscape of sequencing technologies and their impact on microbial genome assemblies and annotation. PLoS ONE 7: 48837.

Mayer, F. L. J., Stalling, D. L., Johnson, J. L. (1972). Phthalate esters as environmental contaminants. Nature 238: 411–413.

McInerney, M., Rohlin, L., Mouttaki, H., Kim, U., Krupp, R. S., Rios-Hernández, L., Sieber, J., Struchtemeyer, C. G., Bhattacharyya, A., Campbell, J. W., Gunsalus, R. P. (2007). The genome of *Syntrophus aciditrophicus*: life at the thermodynamic limit of microbial growth. Proc Natl Acad Sci USA 104: 7600–7605.

Mechichi, T., Stackebrandt, E., Gad’on, N., Fuchs, G. (2002). Phylogenetic and metabolic diversity of bacteria degrading aromatic compounds under denitrifying conditions, and description of *Thauera phenylacetica* sp. nov., Thauera aminoaromatica sp. nov., and Azoarcus buckelii sp. nov. Arch Microbiol 178: 26–35.

Meganathan, R. (2001). Ubiquinone biosynthesis in microorganisms. FEMS Microbiol. Lett. 203: 131–139.

Meng, X. Z., Wang, Y., Xiang, N., Chen, L., Liu, Z. G., Wu, B., et al. (2014). Flow of sewage sludge-borne phthalate esters (PAEs) from human release to human intake: implication for risk assessment of sludge applied to soil. Sci Total Environ 476: 242–249.

Molina-Henares, A. J., Krell, T., Eugenia-Guazzaroni, M., Segura, A., Ramos, J. L. (2006). Members of the IclR family of bacterial transcriptional regulators function as activators and/or repressors. FEMS Microbiol Rev 30: 157–186.

Mountfort, D. O., Brulla, W. J., Krumholz, L. R., Bryant, M. P. (1984). *Syntrophus buswellii* gen. nov., sp. nov. : a benzoate catabolizer from methanogenic ecosystems. Int J Syst Bacteriol 34: 216–217.

Müller, J. A., Schink, B. (2000). Initial steps in the fermentation of 3-hydroxybenzoate by *Sporotomaculum hydroxybenzoicum*. Arch Microbiol 173: 288–295.

Naumov A. V., Gafarov A. B., Boronin A. M. (1996). A novel ortho-phthalate utilizer-An acetic acid bacterium *Acetobacter* sp. Microbiology 65: 182–186.

Nawrocki, E. P., Eddy, S. R. (2013). Infernal 1.1: 100-fold faster RNA homology searches. Bioinformatics. 29: 2933–5.

Net, S., Sempere, R., Delmont, A., Paluselli, A., Ouddane, B. (2015). Occurrence, fate, behaviour and ecotoxicological state of phthalates in different environmental matrices. Environ Sci Technol 49: 4019–4035.

Neuhoff, V., Arold, N., Taube, D., Ehrhardt, W. (1988). Improved staining of proteins in polyacrylamide gels including isoelectric focusing gels with clear background at nanogram sensitivity using coomassie brilliant blue G-250 and R-250. Electrophoresis 9: 255–262.

Niazi, J. H., Prasad, D. T., Karegoudar, T. B. (2001). Initial degradation of dimethylphthalate by esterases from *Bacillus* species. FEMS Microbiol Lett 196: 201–205.

Nilsson, C. (1994). Phthalic acid esters used as plastic additives comparisons of toxicological effects. Swedish National Chemicals Inspectorate, Solna.

Nishizawa, T., Tago, K., Oshima, K., Hattori, M., Ishii, S., Otsuka, S., Senoo, K. (2012). Complete genome sequence of the denitrifying and N_2_O-reducing bacterium *Azoarcus* sp. strain KH32C. J Bacteriol 194: 1255.

Nobu, M. K., Narihiro, T., Tamaki, H., Qiu, Y. L., Sekiguchi, Y., Woyke, T., Goodwin, L., Davenport, K. W., Kamagata, Y., Liu, W. T. (2015). The genome of *Syntrophorhabdus aromaticivorans* strain UI provides new insights for syntrophic aromatic compound metabolism and electron flow. Environ Microbiol 17: 4861–4872.

Nomura, Y., Nakagawa, M., Ogawa, N., Harashima, S., Oshima, Y. (1992). Genes in PHT plasmid encoding the initial degradation pathway of phthalate in Pseudomonas putida. J Ferment Bioeng 74: 333–344.

Notredame, C., Higgins, D., Heringa, J. (2000). T-Coffee: A novel method for multiple sequence alignments JMB, 302: 205–217.

Nozawa, T., Maruyama, Y. (1988a). Anaerobic metabolism of phthalate and other aromatic compounds by a denitrifying bacterium. J Bacteriol 170: 5778–5784.

Nozawa, T., Maruyama, Y. (1988b). Denitrification by a soil bacterium with phthalate and other aromatic compouds as substrates. J Bacteriol 170: 2501–2505.

Parales, R. E., Resnick, S. M. (2006). Aromatic ring hydroxylating dioxygenases. In: Ramos,J. L., Levesque, R. C. (Eds.), Pseudomonas. Springer, Netherlands, pp. 287–340.

Park, J. W., Jung, W. S., Park, S. R., Park, B. C., Yoon, Y. J. (2007). Analysis of intracellular short organic acid-coenzymeA esters from actinomycetes using liquid chromatography-electrospray ionization-mass spectrometry. J Mass Spectrom 42: 1136–1147.

Parte, A. C. (2014). LPSN--list of prokaryotic names with standing in nomenclature. Nucleic Acids Res 42: D613–6.

Patil, Y., Junghare, M., Pester, M., Müller, N., Schink, B. (2015). *Anaerobium acetethylicum* gen. nov., sp. nov., a strictly anaerobic, gluconate-fermenting bacterium isolated from a methanogenic bioreactor. Int J Syst Evol Microbiol. 65: 3289–96.

Payne, K. A. P. et al. (2015). New cofactor supports a,b-unsaturated acid decarboxylation via 1,3-dipolar cyclo addition. Nature 14560.

Pérez-Pantoja, D, Gonzaelez1, B. Pieperi, D. H. (2009). “Aerobic degradation of aromatic hydrocarbons” in handbook of hydrocarbon and lipid microbiology, ed. K. N. Timmis (Berlin: Springer-Verlag), p. 799–837.

Peijnenburg, W. J. G. M., Struijs, J. (2006). Occurrence of phthalate esters in the environment of the Netherlands. Ecotoxicol Environ Saf 63: 204–215.

Perkins, D. N., Pappin, D. J. C., Creasy, D. M., Cottrell, J. S. (1999). Probability-based protein identification by searching sequence databases using mass spectrometry data. Electrophoresis 20: 3551–3567.

Peters, F., Heintz, D., Johannes, J., Dorsselaer, A.V., Boll, M. (2007). Genes, enzymes, and regulation of para cresol metabolism in *Geobacter metallireducens*. J.Bacteriol 189: 4729–4738.

Peters, F., Rother, M., Boll, M. (2004). Selenocysteine-containing proteins in anaerobic benzoate metabolism of *Desulfococcus multivorans*. J Bacteriol 186: 2156–2163.

Pettersen, E. F., Goddard, T. D., Huang, C. C., Couch, G. S., Greenblatt, D. M., Meng, E. C., Ferrin, T. E. J. (2004). UCSF Chimera--a visualization system for exploratory research and analysis. Comput Chem. 25: 1605–12.

Pfennig, N. (1978). *Rhodocyclus purpureus* gen. nov. sp. nov., a ring-shaped, vitamin B12-requiring member of the family *Rhodospirillaceae*. Int J Syst Bacteriol 28: 283–288.

Pfennig, N., Wagener, S. (1986). An improved method of preparing wet mounts for photomicrographs of microorganisms. J Microbiol Meth 4: 303–306.

Philipp, B., Schink, B. (2012). Different strategies in anaerobic biodegradation of aromatic compounds: nitrate reducers versus strict anaerobes. Environ Microbiol Rep 4: 469–478.

Plugge, C. M., Zhang, W., Scholten, J. C. M., Stams, A. J. M. (2011). Metabolic flexibility of sulfate-reducing bacteria. Front. Microbiol 2, 81.

Pruesse, E., Peplies, J. & Glöckner, F. O. (2012). SINA: Accurate high-throughput multiple sequence alignment of ribosomal RNA genes. Bioinformatics 28: 1823–1829.

Porebski, S., Bailey, L., Baum, B. (1997). Modification of a CTAB DNA extraction protocol for plants containing high polysaccharide and polyphenol components. Plant Mol Biol Rep 15: 8–15.

Powlowski, J., Shingler, V. (1994). Genetics and biochemistry of phenol biodegradation by *Pseudomonas* sp. CF600. Biodegradation 5: 219–236.

Pruesse, E., Quast, C., Knittel, K., Fuchs, B. M., Ludwig, W., Peplies, J., Glöckner, F. O. (2007). SILVA: a comprehen-sive online resource for quality checked and aligned ribosomal RNA sequence data compatible with ARB. Nucleic Acids Res 35: 7188–7196.

Pujar, B. G., Ribbons, D. W. (1985). Phthalate Metabolism in *Pseudomonas fluorescens* PHK: purification and properties of 4, 5-dihydroxyphthalate decarboxylase. Appl Environ Microbiol 49: 374–376.

Qiu, Y. L., Sekiguchi, Y., Hanada, S., Imachi, H., Tseng, I. C., Cheng, S. S., Ohashi, A., Harada, H., Kamagata, Y. (2006). *Pelotomaculum terephthalicum* sp. nov. and *Pelotomaculum isophthalicum* sp. nov.: two anaerobic bacteria that degrade phthalate isomers in syntrophic association with hydrogenotrophic methanogens. Archives Arch Microbiol 185: 172–182.

Qiu, Y. L., Sekiguchi, Y., Imachi, H., Kamagata, Y., Tseng, I. C., Cheng, S. S., Ohashi, A., Harada, H. (2004). Identification and isolation of anaerobic, syntrophic phthalate isomer-degrading microbes from methanogenic sludges treating wastewater from terephthalate manufacturing. Appl Environ Microbiol 70: 1617–1626.

Quast, C., Pruesse, E., Yilmaz, P., Gerken, J., Schweer, T., Yarza, P., Peplies, J., Glöckner, F. O. (2013). The SILVA ribosomal RNA gene database project: improved data processing and web-based tools. Nucl Acids Res 41: 590–596.

Rabus, R., Kube, M., Heider, J., Beck, A., Heitmann, K., et al. (2005). The genome sequence of an anaerobic aromatic-degrading denitrifying bacterium, strain EbN1. Arch Microbiol 183: 27–36.

Rangarajan, E. S. et al. (2004). Crystal structure of a dodecameric FMN-dependent UbiXlike decarboxylate (Pad1) from *Eschericia coli* O157:H7. Protein Sci 13: 3006–3016.

Rather, L. J. et al. (2011). Structure and mechanism of the diiron benzoyl-coenzyme A epoxidase BoxB. J. Biol. Chem. 286, 29241–29248.

Reinhold, H. B., Hurek, T., Gillis, M., Hoste, B., Vancanneyt, M., Kersters, K., De, L. J. (1993). *Azoarcus* gen. nov., nitrogen-fixing *Proteobacteria* associated with roots of kallar grass (*Leptochloa fusca* (L.) Kunth), and description of two species, Azoarcus indigens sp. nov. and Azoarcus communis sp. nov. Int J Syst Bacteriol 43: 574–584.

Ribbons, D. W., Eaton, R. W. (1982). Chemical transformations of aromatic hydrocarbons that support the growth of microorganisms. A. M. Chakrabarty. CRC Press, Boca Raton, Fla. In: Biodegradation and detoxification of environmental pollutants (Ed.) 59–84.

Ribbons, D. W., Keyser, P., Kunz, D. A., Taylor, B. F. (1984). Microbial degradation of phthalates. In: Gibson, D.T. (Ed.), Microbial degradation of organic compounds Marcel Dekker, New York.

Robert, X., Gouet, P., (2014). Deciphering key features in protein structures with the new ENDscript server’. Nucl. Acids Res. 42: W320–W324.

Saitou, N., Nei, M. (1987). The neighbor-joining method: a new method for reconstructing phylogenetic trees. Mol Biol Evol 4: 406–425.

Sambrook, J., Fritsch, E. F., Maniatis, T. (1989). Molecular cloning: a laboratory manual, 2nd edition. Cold Spring Harbor Laboratory, Cold Spring Harbor, New York.

Schink, B. (1997). Energetics of syntrophic cooperation in methanogenic degradation. Microbiol Mol Biol Rev 61: 262–280.

Schink, B., Brune, A., Schnell, S. (1992). Anaerobic degradation of aromatic compounds. In Microbial degradation of natural compounds. Winkelmann, G. (ed.). Weinheim: VCH, p. 219–242.

Schink, B., Philipp, B., Müller, J. (2000). Anaerobic degradation of phenolic compounds. Natur wissenschaften 87: 12–23.

Schläfli, H. R., Weiss, M. A., Leisinger, T. (1994). Terephthalate 1, 2-dioxygenase system from *Comamonas testosteroni* T-2: purification and some properties of the oxygenase component. J Bacteriol 176: 6644–6652.

Schmidt, A., Müller, N., Schink, B., Schleheck, D. (2013). A proteomic view at the biochemistry of syntrophic butyrate oxidation in *Syntrophomonas wolfei*. PLoS ONE 8: 56905.

Schöcke, L., Schink, B. (1999). Energetics and biochemistry of fermentative benzoate degradation by *Syntrophus gentianae*. Arch Microbiol 171: 331–337.

Schwarzbauer, J., Heim, S., Brinker, S., Littke, R. (2002). Occurrence and alteration of organic contaminants in seepage and leakage water from a waste deposit land fill. Water Res 36: 2275–2287.

Sepic, E., Bricelj, M., Leskovsek, H. (1998). Degradation of fluoranthene by *Pasteurella* sp. IFA and Mycobacterium sp. PYR-1, Isolation and identification of metabolites. J Appl Microbiol 85: 746–754.

Shigematsu, T., Yumihara, K., Ueda, Y., Morimura, S., Kida, K. (2003). Purification and gene cloning of the oxygenase component of the terephthalate 1,2-dioxygenase system from *Delftia tsuruhatensis* strain T7. FEMS Microbiol Lett 220: 255–260.

Sin, J. C., Lam, S. M., Mohamed, A. R., Lee, K. T. (2012). Degrading endocrine disrupting chemicals from wastewater by TiO2 photocatalysis: A Review. International Journal of Photoenergy p. 23.

Stamatakis, A., Hoover, P., Rougemont, J. (2008). A rapid bootstrap algorithm for the RAxML web servers. Syst Biol 57: 758–771.

Staples, C. A., Peterson, D. R., Parkerton, T. F., Adams, W. J. (2002). The environmental fate of phthalate esters: a literature review. Chemosphere 35: 667–749.

Szewzyk, R., Pfennig, N. (1987). Complete oxidation of catechol by the strictly anaerobic sulfate-reducing *Desulfobacterium catecholicum* sp., nov. Arch Microbiol 147: 163–168.

Szewzyk, U., Schink, B. (1989). Degradation of hydroquinone, gentisate, and benzoate by a fermenting bacterium in pure or defined mixed culture. Arch Microbiol 151: 541–545.

Tamura, K., Dudley, J., Nei, M., Kumar, S. (2007). MEGA4: molecular evolutionary genetics analysis (MEGA) software version 4.0. Mol Bio Evol 24: 1596–1599.

Tamura, K., Peterson, D., Peterson, N., Stecher, G., Nei, M., Kumar, S. (2011). MEGA5: Molecular evolutionary genetics analysis using Maximum Likelihood, Evolutionary Distance, and Maximum Parsimony Methods. Mol Biol Evol 28: 2731–2739.

Tartoff, K. D., Hobbs, C. A. (1987). Improved media for growing plasmid and cosmid clones. Bethesda Res Lab Focus 9: 12.

Tasaki, M., Kamagata, Y., Nakamura, K., Mikami, E. (1991). Isolation and characterization of a thermophilic benzoate-degrading, sulfate-reducing bacterium, Desulfotomaculum thermobenzoicum sp. nov. Arch Microbiol 155: 348–352.

Taylor, B. F., Ribbons, D. W. (1983). Bacterial decarboxylation of o-phthalic acids. Appl Environ Microbiol 46: 1276–1281.

Unciuleac, M., Boll, M. (2001). Mechanism of ATP-driven electron transfer catalyzed by the benzene ring-reducing enzyme benzoyl-CoA reductase. Proc Natl Acad Sci USA 98: 13619–13624.

Vaillancourt, F. H., Bolin, J. T., Eltis, L. D. (2006). The ins and outs of ring cleaving dioxygenases. Crit Rev Biochem Mol Biol 41: 241–267.

Vamsee-Krishna, C., Mohan, Y., Phale P. S. (2006). Biodegradation of phthalate isomers by *Pseudomonas aeruginosa* PP4, *Pseudomonas* sp. PPD and *Acinetobacter lwoffii* ISP4. Appl Microbiol Biotechnol 72: 1263–1269.

Vats, S., Singh, K. R., Tyagi, P. (2013). Phthalates-a priority pollutant. I J A B R 3: 1–8.

Wang, Y. P., Gu, J. D. (2006a). Degradability of dimethyl terephthalate by *Variovorax paradoxus* T4 and *Sphingomonas yanoikuyae* DOS01 isolated from deep-ocean sediments. Ecotoxicology 15: 549–557.

Wang, Y. P., Gu, J. D. (2006b). Degradation of dimethyl iso-phthalate by *Viarovorax paradoxus* strain T4 isolated from deep-ocean sediment of the south china sea. Human Ecol Risk Assess 12: 236–247.

Wang, Y. Y., Fan, Y. Z., Gu, J. D. (2003). Aerobic degradation of phthalic acid by *Comamonas acidovoran* Fy-1 and dimethyl phthalate ester by two reconstituted consortia from sewage sludge at high concentrations. World J Microbiol Biotechnol 19: 811–815.

Wang, Y. Z., Zhou, Y., Zylstra, G. J. (1995). Molecular analysis of isophthalate and terephthalate degradation by *Comamonas testosteroni* YZW-D. Environ Health Persp 103: 9–12.

Weelink, S. A. B., van Eekert, M. H. A, Stams, A. J. M. (2010). Degradation of BTEX by anaerobic bacteria: physiology and application. Rev Environ Sci Biotechnol 9: 359–385.

Wensing, M., Uhde, E., Salthammer, T. (2005). Plastics additives in the indoor environment-flame retardants and plasticizers. Sci Total Environ 339: 19–40. White, M. D., Payne, K. A. P., Fisher, K., Marshall, S. A., Parker, D., Rattray, N. J. W., Trivedi, D. K., Goodacre, R., Rigby, S. E. J., Scrutton, N. S., Hay, S., Leys, D. (2015). UbiX is a flavin prenyltransferase required for bacterial ubiquinone biosynthesis. Nature 522: 502–506.

White, M. D., Payne, K. A. P., Fisher, K., Marshall, S. A., Parker, D., Rattray, N. J. W., et al. (2015). UbiX is a flavin prenyltransferase required for bacterial ubiquinone biosynthesis. Nature 522: 502–506.

Widdel, F., Bak, F. (1992). Gram negative mesophilic sulfate reducing bacteria. In: Balows H, Truper HG, Dworkin M, Harder W, Schleifer KH (eds) The Prokaryotes Vol. IV. Chapter 183: 3352–3378. New York, Berlin, Heidelberg: Springer.

Widdel, F., Kohring, G. W., Mayer F. (1983). Studies on dissimilatory sulfate-reducing bacteria that decompose fatty acids. Characterization of the filamentous gliding *Desulfonema limicola*. Arch Microbiol 134: 286–294.

Widdel, F., Schnell, S., Heising, S., Ehrenreich, A., Abmus, B., Schink, B. (1993). Anaerobic ferrous iron oxidation by anoxygenic phototrophs. Nature 362: 834–836.

Wischgoll, S., Heintz, D., Peters, F., Erxleben, A., Sarnighausen, E., Reski, R., Dorssalaer, A. V., Boll, M. (2005). Gene clusters involved in anaerobic benzoate degradation of *Geobacter metallireducens*. Mol Microbiol 58: 1238–1252.

Woodward, K. N. (1988). The adipates in phthalate esters: toxicity and metabolism. vol. II. pp. 168-173, Boca Raton, FL. CRC press.

Woodward, K. N. (1990). Phthalate esters, cystic kidney disease in animals and possible effects on human health: a review. Hum Exp Toxicol 9: 397–401.

Wu, X. L., Liang, R. X., Dai, Q. Y., Jin, D. C., Wang, Y. Y., Chao, W. L. (2010). Complete degradation of di-n-octyl phthalate by biochemical cooperation between *Gordonia* sp. strain JDC-2 and *Arthrobacter* sp. strain JDC-32 isolated from activated sludge. J Hazard Mater 176: 262–268.

Zhang, H., Javor, G. T. (2000). Identification of the ubiD gene on the *Escherichia coli* chromosome. J Bacteriol 182: 6243–6246.

Zheng, Z., He, P. J., Shao, L. M., Lee, D. J. (2007). Phthalic acid esters in dissolved fractions of land fill leachates. Water Res 41: 4696–4702.

Zuckerkandl, E., Pauling, L. (1965). Evolutionary divergence and convergence in proteins. edited in evolving genes and proteins by V. Bryson and H.J. Vogel, pp. 97–166. Academic Press, New York.

